# The lung employs an intrinsic surfactant-mediated inflammatory response for viral defense

**DOI:** 10.1101/2023.01.26.525578

**Authors:** Sandra L. Leibel, Rachael N. McVicar, Rabi Murad, Elizabeth M. Kwong, Alex E. Clark, Asuka Alvarado, Bethany A. Grimmig, Ruslan Nuryyev, Randee E. Young, Jamie Casey Lee, Weiqi Peng, Yanfang Peipei Zhu, Eric Griffis, Cameron J. Nowell, Kang Liu, Brian James, Suzie Alarcon, Atul Malhotra, Linden J. Gearing, Paul J. Hertzog, Cheska Marie Galapate, Koen M.O. Galenkamp, Cosimo Commisso, Davey M. Smith, Xin Sun, Aaron F. Carlin, Ben A. Croker, Evan Y. Snyder

**Affiliations:** Department of Pediatrics, University of California San Diego School of Medicine; La Jolla, CA, 92093, USA; Sanford Consortium for Regenerative Medicine; La Jolla, CA, 92037, USA; Sanford Burnham Prebys Medical Discovery Institute, Center for Stem Cells & Regenerative Medicine; La Jolla, CA, 92037, USA; Department of Medicine, University of California San Diego School of Medicine; La Jolla, CA, 92093, USA; Nikon Imaging Center, University of California San Diego School of Medicine; La Jolla, CA, 92093, USA; Monash Institute of Pharmaceutical Sciences; Parkville, Victoria, 3052, Australia; La Jolla Institute for Immunology; La Jolla, CA, 92037, USA; Division of Pulmonary, Critical Care, and Sleep Medicine, University of California San Diego; CA, 92093, USA; Centre for Innate Immunity and Infectious Diseases, Hudson Institute of Medical Research; Clayton, Victoria, 3168, Australia; Department of Molecular and Translational Sciences, Monash University; Clayton, Victoria, 3168, Australia; Sanford Burnham Prebys Medical Discovery Institute, Cell and Molecular Biology of Cancer Program; La Jolla, CA, 92037, USA

## Abstract

Severe Acute Respiratory Syndrome Coronavirus-2 (SARS-CoV-2) causes an acute respiratory distress syndrome (ARDS) that resembles surfactant deficient RDS. Using a novel multi-cell type, human induced pluripotent stem cell (hiPSC)-derived lung organoid (LO) system, validated against primary lung cells, we found that inflammatory cytokine/chemokine production and interferon (IFN) responses are dynamically regulated autonomously within the lung following SARS-CoV-2 infection, an intrinsic defense mechanism mediated by surfactant proteins (SP). Single cell RNA sequencing revealed broad infectability of most lung cell types through canonical (ACE2) and non-canonical (endocytotic) viral entry routes. SARS-CoV-2 triggers rapid apoptosis, impairing viral dissemination. In the absence of surfactant protein B (SP-B), resistance to infection was impaired and cytokine/chemokine production and IFN responses were modulated. Exogenous surfactant, recombinant SP-B, or genomic correction of the SP-B deletion restored resistance to SARS-CoV-2 and improved viability.

## INTRODUCTION

Despite extensive study over the past 2 years, our understanding of pulmonary coronavirus disease-19 (COVID-19) caused by infection with Severe Acute Respiratory Syndrome Coronavirus-2 (SARS-CoV-2), remains incomplete and ever-shifting as new viral variants emerge; consequently, treatment recommendations vacillate (*1–3*). We found particularly striking – and informative – the evolution in thinking about the nature of the acute respiratory distress syndrome seen in COVID-19 patients (CARDS) (*4*). Autopsy material from patients who succumbed to CARDS showed a reduction in surfactant protein (SP) gene expression in the lung (*5, 6*).

Surfactants are complexes secreted by pulmonary alveolar type 2 (AT2) cells that reduce surface tension in the alveoli to facilitate gas exchange (*7, 8*) and by club cells to maintain the airway epithelial lining fluid (*9*). Expression of surfactant protein B (SP-B), the most critical component of surfactant, is gestationally regulated (*10, 11*), and a developmental deficiency of surfactant is the most common cause of neonatal respiratory distress syndrome (RDS) in preterm newborns (*12*). Such RDS is characterized by poor pulmonary compliance and refractory hypoxemia (*13, 14*). It was intriguing that CARDS might share a similar pathophysiology with that early developmental form of RDS (*15, 16*) although more evidence is needed. Additionally, exogenous pulmonary surfactant (a therapy for neonatal RDS) has also been proposed as a therapy to restore lung function following COVID-19 (*17, 18*).

We studied the response to infection of multiple variants of SARS-CoV-2 in a unique *in vitro* system that reproduces the development and three-dimensional (3D) cytoarchitecture of most cell types of the lung. We derived 3D lung organoids (LOs) representing the proximal and distal portions of the lung, including mesenchymal, goblet, serous, basal, pulmonary neuroendocrine, club, alveolar type 1, and surfactant-producing AT2 cells (*19–22*). Most of these cell types were infectable with SARS-CoV-2 (*23–26*) – surprising given the wide acceptance of the virus’ canonical routes of entry. Equally surprising was learning, through transcriptional analysis of the downstream effects of SARS-CoV-2 infection, that surfactant plays a critical role in the lung’s defense mechanisms against acute viral infection, likely defined by a heretofore unrecognized intrinsic intra-pulmonary inflammatory feedback loop that autonomously attempts to restore lung homeostasis after infection by inhibiting viral dissemination, averting lung epithelial demise, and dampening inflammatory cascades. This process has not previously been described and may offer avenues for new COVID-19 treatments and novel therapeutic strategies against other viruses.

## RESULTS

### Human induced pluripotent stem cell (hiPSC)-derived lung organoids (LOs) are susceptible to infection with multiple strains of SARS-CoV-2, validated in primary human lung epithelia

Previous studies of SARS-CoV-2 have relied primarily on post-mortem material (*5, 6, 27, 28*), lung-derived cell lines (often neoplastic) (*29, 30*), and on cultures of selected lung constituents (*26*). We employed a system that enabled us to map -- prospectively and in an unbiased manner -- the full downstream transcriptional, proteomic, and functional response of human pulmonary epithelial and mesenchymal cells of the normal lung upon first exposure to variants of SARS-CoV-2. This system was derived from an optimized patient-specific culture system that contains epithelial and mesenchymal functional lung components in a 3D cytoarchitecture, as previously devised and further refined for this study (*19, 21, 22*).

Creating these lung organoids (LOs) emulate normal lung development by exposing human induced pluripotent stem cells (hiPSCs), which model the epiblast, to a precise sequence of inductive factors **[Fig. 1A and Supplemental Figure 1a]** (see **Methods** for details). Briefly, we generated definitive endoderm from hiPSCs through Nodal signaling with high dose Activin A (*31*). Anterior foregut endoderm (AFE) was generated following inhibition of BMP, WNT, and TGFß signaling (*32*). Lung progenitor cells (LPCs) were then ventralized from AFE (*33, 34*) via passaging AFE cells in matrigel (*35*) while modulating BMP, WNT, and retinoic acid (RA) signaling. *NKX2-1* was upregulated in early LPCs in the ventral aspect of the AFE to regulate lung development and surfactant activation (*36*). The LPCs were further differentiated into 3 distinct heterogeneous LOs made up of lung cells located in the proximal or distal areas of the human lung. Whole lung organoids (WLO) represented many of the cell types of the lung, including airway cells, alveolar cells and PNECs, and were induced via the GSK3ß inhibitor CHIR99021, FGF7, FGF10, EGF, RA, VEGF/PGF, “*<u>d</u>*examethasone, *<u>c</u>*AMP, and *i*sobutylxanthine” (“DCI”) in a stepwise manner, beginning from LO induction, to branching morphogenesis, and finally, maturation (*19, 20*). Exposure to DCI resulted in phenotypic maturation of the organoids within 24 hours of exposure, “ballooning” in response to developmentally appropriate induction **[Supplementary Movie 1]**, assuming a structure reminiscent of a “mini-lung” **[Supplemental Fig. 1B]**. These structures contained ATII cells (pro-SPC**+**) that produced surfactant – both anatomically and by protein expression (*20*) – as well as alveolar type I (ATI) cells (HT1-56**+**), goblet, club, and basal cells **[Supplementary Fig. 1C]**. Importantly, surfactant-producing ATII cells developed only when the hiPSC-derivatives were permitted to assume a 3D configuration (*20*). Mesenchymal cells, including smooth muscle, were present **[Supplementary Fig. 1C]**. By altering growth factor exposure to the LPCs, including FGF2, high dose FGF10, and DCI, we generated proximal lung organoids (PLO) **[Supplementary Fig. 1B]** which contained more airway cells (*22*) including basal (p63**+**), club (SCGB3A2**+**), goblet, and ciliated cells (Ac-Tub**+**) **[Supplementary Fig. 1D, Supplementary Movie 2].** Distal lung organoids (DLO) were derived via exposure of LPCs to CHIR99021, SB431542, FGF7, RA and DCI (*21*) **[Supplementary Fig. 1B, D]**. These had the highest expression of alveolar cells.

**FIGURE 1:**
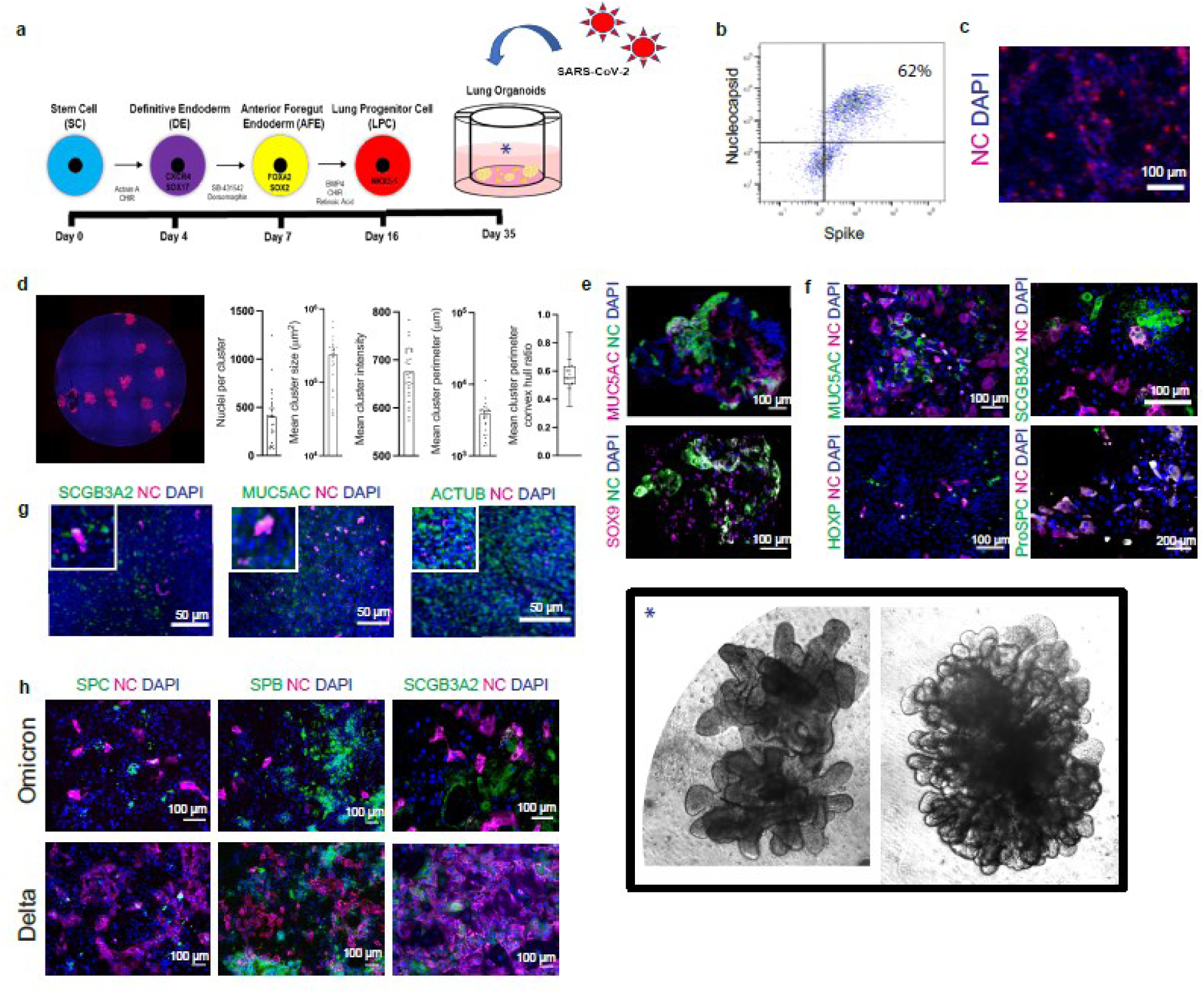
Human induced pluripotent stem cell (hiPSC)-derived lung organoids (LOs) are susceptible to infection with multiple strains of SARS-CoV-2, validated in primary human lung epithelia. **(a)** Schematic of iterative steps by which hiPSCs are instructed to become first definitive endoderm, then anterior foregut endoderm, then lung progenitors cells, and finally the array of multiple pulmonary epithelial cell types that assemble into a 3D structure with a form and cytoarchitecture that resembles a lung in situ. See **Supplementary Fig. 1** for greater detail. Photomicrographs of actual whole lung organoids (WLOs) (drawn in the schematic [**asterisk \***]) are magnified in the **inset [*]** in the lower right of the montage. These WLOs and/or specialized portions of the lung (e.g., proximal lung organoids [PLOs] or distal lung organoids [DLOs]) were cultured in serum free basal medium and infected with a broad range of SARS-CoV-2 viral variants (**red multi-spiked circles** in the schematic) for 24-72 hours. **(b)** Representative flow cytometry of infected 3D PLOs expressing viral proteins Spike and Nucleocapsid (NC) confirms that 62% of the pulmonary epithelial cells were infected. **(c)** Immunostaining of infected dissociated PLOs at 24 hpi for viral NC **(red)** confirms the cytometric findings of viral infection. **(d)** Immunostaining of infected dissociated PLOs overlayed with carboxymethycellulose to enhance visualization and permit quantification. Cells were fixed, permeabilized, and co-stained with NC-AF594 antibody **(red)** +DAPI **(blue)**. A representative well is shown. Images were quantified with a custom-scripted Image J code. Measurements of NC**^+^** nuclei/cluster, cluster size and perimeter, and cluster intensity confirm robust infection. In some experiments, LOs were dissociated acutely to enable unambiguous visualization of individual cells without the risk of one cell being superimposed upon another. **(e)** Immunostaining of *intact* 3D PLOs 36 hpi for co-expression of viral NC (indicative of infection) in multiple pulmonary epithelial cell types as identified by cell type-specific immunomarkers. Two representative cell types are shown: goblet cells (MUC5AC) and lung progenitor cells (SOX9). **(f)** Immunostaining of dissociated 3D LOs with viral NC and cell-type lung epithelial markers: goblet cells (MUC5AC), club cells (SCGB3A2), alveolar type 2 cells (proSPC), alveolar type 1 cells (HOXP). **(g)** To validate the range of cell types infected in LOs, primary human bronchial epithelial cell (HBEC) air liquid interphase (ALI) cultures were also infected and immunostained 24 hpi. The results in these primary cells confirmed the findings in the LOs. Shown in the co-expression of viral NC (evidence of infection) in a range of the same representative pulmonary epithelial cells show above: goblet cells (MUC5AC), club cells (SCGB3A2), ciliated cells (AcTub). **Insets** are 2.5 x original images. **(h)** Immunostaining of dissociated LOs, 24 hpi, comparing SARS-CoV-2 variants Omicron vs. Delta. The data are representative of at least three independent experiments. The lung immunomarkers identify the cells described above. SPB and SPC both identify alveolar type 2 cells. In all photomicrographs, a DAPI nuclear stain (**blue**) is used to visualize all cells in the field. All scale bar = 100 µm unless otherwise specified.

Single cell RNA sequencing (scRNAseq) profiling confirmed the expression of lung epithelial and mesenchymal cell markers in each of the LO types **[Supplementary Figure 2A-C]**, as determined by correlating the signature genes to single cell sequencing data of published data sets from fetal and adult human lung cells (*37–44*) and the gene annotation and analysis resource, Metascape (*45*). In some cases, to address the issues of COVID-19 disease disparity, we generated patient-specific LOs using hiPSC from men and women of different racial and ethnic backgrounds (e.g., African American, Caucasian, Hispanic) **[Supplementary Table 1]**.

To determine the acute and primary tropism of the SARS-CoV-2 spike protein in the lung without permitting intercellular spread, which can confound discerning the initial target, we first infected the LOs with a replication-incompetent vesicular stomatitis virus (VSV) pseudotyped with the SARS-CoV-2 spike protein conjugated to GFP (pseudovirus-GFP) (*46*) **[Supplementary Fig. 3A]** at a multiplicity of infection (MOI) of 1-5. Single cell RNAseq of the infected samples, after sorting for GFP**+** (infected) and GFP**–** (uninfected) cells showed differing clusters in the UMAP **[Supplementary Fig. 3B]**. The basal cells and differentiating epithelial cell genes were highly represented in the pseudoviral-infected GFP+ samples **[Supplementary Fig. 3B, 3C]**. To confirm the cell types infected, we performed immunofluorescence staining of acutely dissociated LOs in monolayer (to enhance visualization of individual cells), 24 hours post infection (hpi), with antibodies against P63 (basal cell), SCGB3A2 (secretory cell), and MUC5AC (goblet cell) **[Supplementary Fig. 3D]**. As expected, there was co-localization of pseudovirus-GFP with MUC5AC**+** and SCGB3A2**+** cells. Co-localization of pseudovirus-GFP with MUC5AC**+** was also detected in the intact 3D lung organoids **[Supplementary Fig. 3E]**.

Next, we infected the 3D proximal LOs with authentic replication-competent SARS-CoV-2 virus, variant WA1 (USA/WA1-2020, A) at an MOI of 5. Infection was confirmed in all LOs by flow cytometry using antibodies specific for the viral nucleocapsid (NC) and spike protein (S). **[Fig. 1B]**. The 3D LOs were dissociated, plated as a monolayer, then infected with 150 PFU of WA1 for 24 hours **[Fig. 1C]** or infected for one hour and then overlayed with carboxymethycellulose for 24 hours and infected cells (NC antibody **+**) were quantified with a custom-scripted Image J code **[Fig. 1D]**. To determine the lung cells susceptible to infection, we stained intact and dissociated LOs for specific lung markers, and the viral NC. In the 3D WLOs, MUC5AC**+** (goblet) and SOX9**+** (progenitor) cells were infected **[Fig. 1E]**, and in the dissociated PLOs and DLOs, SCGB3A2**+** (secretory), MUC5AC**+** (goblet) and pro-SPC**+** (ATII) cells were infected, but not HOPX**+** (ATI) cells **[Fig. 1F]**. The tropism of the virus for these cell types in the organoids was validated in primary human lung tissue: in HBEC-derived airway epithelial cells, SARS-CoV-2 similarly infected SCGB3A2**+** (secretory), AcTub**+** (ciliated), and MUC5AC**+** (goblet) cells **[Fig. 1G]**.

To explore the differing pathology of SARS-CoV-2 variants, we infected PLOs and DLOs with the Delta (hCoV-19/USA/PHC658/2021, B.1.617.2, Delta) and Omicron (hCoV-19/USA/CA-SEARCH-59467/2021, BA.1.20, Omicron) variants and examined cellular targets and viral dissemination. As found above for WA1, the Delta and Omicron variants infected a similar spectrum of pulmonary cells **[Fig. 1H]**. The Delta variant disseminated in culture more efficiently in 24h than the Omicron variant. The Delta variant infected SPC+ ATII cells in the DLOs, whereas Omicron did not, suggesting that the tropism of the Omicron variant may not be in distal cells compared to other variants of SARS-CoV-2. This observation is consistent with clinical evidence that Delta causes a more distal lung pathology whereas Omicron causes more upper airway inflammation.

### Transcriptional changes and tropism in hiPSC-LOs infected with SARS-CoV-2; most cell types in lung organoids (LOs) can be infected by SARS-CoV-2, independent of ACE2

To gain insight into the response of lung epithelial cells to SARS-CoV-2, we performed unbiased genome-wide transcriptome profiling on proximal lung organoids (PLOs) at 0 (mock), 3, and 24 hpi with the Delta variant. Six hundred and thirty-seven differentially expressed genes (DEGs) were identified between 0 and 3 hpi, and 1302 DEGs were identified between 0 and 24 hpi. Principal component analysis (PCA) demonstrated that PLOs at 24 hpi occupied a distinct transcriptional space compared to mock-infected PLOs **[Fig. 2a]**. Volcano plots of the mock-infected vs. 24 hpi PLOs showed induction of multiple chemokine and cytokine transcripts including *CCL22; CXCL-1, -2, -3, -5* and *-6; IFNE;* and *CSF3* (the latter being the most upregulated in infected HBECs post-infection (*47*)) **[Fig. 2b]** Interestingly, *HOPX*, a gene associated with alveolar cells and which functions to restore epithelial barriers and prevent inflammatory cell infiltration (*48, 49*), was downregulated but was not a primarily infected cell type **[Fig. 1f]**. *TXNRD1*, an enzyme important in protecting cells against oxidative stress (*50*) was also decreased.

**FIGURE 2:**
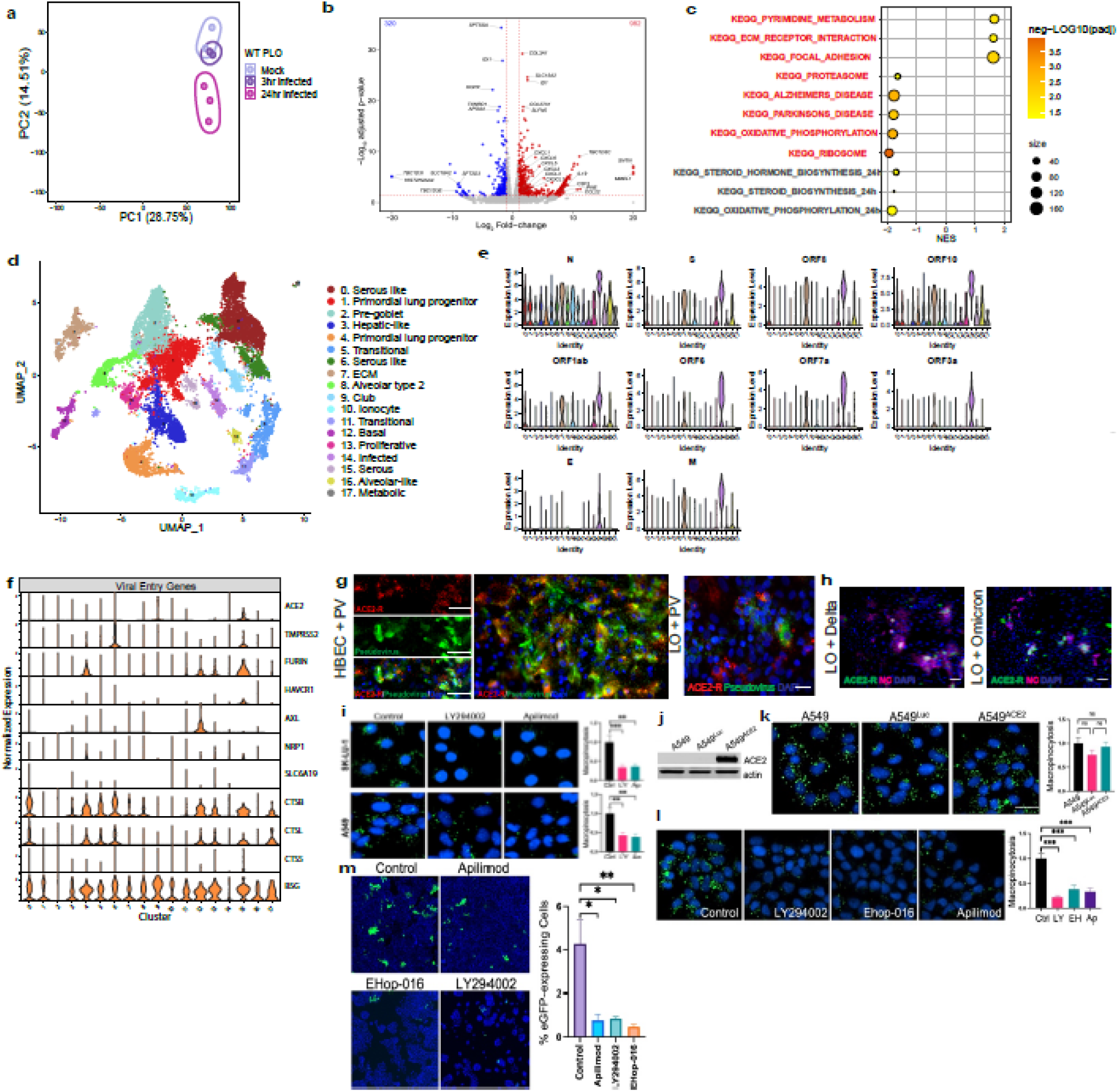
Transcriptional changes and tropism in hiPSC-LOs infected with SARS-CoV-2; most cell types in lung organoids (LOs) can be infected by SARS-CoV-2, independent of ACE2. **(a)** Principal component analysis (PCA) plot of the WT proximal lung organoids (PLO) comparing samples that were uninfected (mock), 3 hpi and 24 hpi. Note that PLOs infected for 24 hours segregate from those infected for only 3 hours or mock-infected. **(b)** Volcano plot analysis of differential expression of SARS-CoV-2 infected PLOs versus mock infection. The red lines indicate a p-adjusted value <0.05. Note a heavy upregulation (red dots) of inflammatory cytokines within the PLOs, even without the addition of hematopoietic or vascular derivatives from the hiPSCs or perfusion of the organoids. **(c)** Gene over-representation analysis using Kyoto Encyclopedia of Genes and Genomes (KEGG) pathway database of SARS-CoV-2 3 hpi (red text) and 24 hpi (black text) vs mock infected LOs. **(d)** UMAP of 3D PLOs, 24 hpi with the alpha variant. Clusters were generated via Harmony Integration of 3 independent samples. ECM = extracellular matrix. **(e)** SARS-CoV-2 viral genes in each cluster as violin plots. The vertical bars represent the expression level. Note that every cluster expresses viral genes. **(f)** Violin plots of SARS-CoV-2 entry genes in each cluster in the PLOs. Again, note that such genes are present in every cluster. **(g)** Immunofluorescence images of primary human bronchial epithelial cells (HBEC) subjected to air-liquid interface (ALI) methodology (left panel) (serving as a primary lung cell control) and hiPSC-derived PLOs (acutely dissociated to allow excellent exposure to the virus and visualization without overlapping cells) (right panel) infected with SARS-CoV-2 pseudovirus (to allow examination of viral entry without other disruption or killing of the host cell). Images show presence of pseudovirus in cells with and without the ACE2 receptor. Furthermore, not all ACE2 immunopositive cells are infected. **(h)** Immunofluorescence images of dissociated hiPSC-PLOs infected with Delta and Omicron variants. Images show presence of NC in cells with and without the ACE2 receptor. **(i)** Proof that lung cells employ an endocytotic mechanism for intracellularly entry (entrance of green fluorescent dextran) that Apilimod can specifically block, making that drug a useful tool for identifying endocytosis as a cell entry mechanism. Shown is endocytosis (via macropinocytosis) by FITC-Dextran uptake. As positive controls, SK-LU-1 lung cells were treated with 25 µM LY294002 (PI3K inhibitor) and A549 lung cells were treated with 100 µM LY294002 – both known to block endocytosis, and against which Apilimod was compared. Apilimod (phosphatidylinositol-3-phosphate 5-kinase (PIKfyve) inhibitor) was used at 100 nM for both cell lines. Cells were pre-treated with the drugs for 3 hours prior to FITC-Dextran incubation for 30 min. Vehicle (DMSO) was used as a control. Endocytosis/macropinocytosis (green fluorescence) was inhibited comparably and significantly by all drugs. **p<0.01, ***p<0.001. **(j)** To show that ACE2 receptor expression in lung cells does not affect the extent of endocytosis, A549 lung cells were engineered to express ACE2 or luciferase (Luc) as a control. Actin was used as a loading control in the western blot. **(k)** FITC-Dextran uptake showing intact endocytosis in A549 wild type, A549^Luc^ and A549^ACE2^ cells. There was no significant difference in endocytosis/macropinocytosis. ns=not significant. **(l)** A549^ACE2^ cells treated with LY294002, 20 mM of Ehop-016 (another known inhibitor of endocytosis via its suppression of Rac1 which perturbs actin cytoskeleton dynamics, and hence another positive control); (c) Apilimod; or (d) control (DMSO), Apilimod or control (DMSO). Apilimod and LY294002 treatment was for 3 hours, while Ehop-016 was used for 2 hours. Endocytosis in A549^ACE2^ cells was still inhibited significantly by LY294002, Apilimod and Ehop-016. ***p<0.001 **(m)** SARS-CoV-2-pseudovirus mediated infection in lung cells, both expressing and not expressing ACE2 receptor, after pre-treatment with LY294002, Apilimod, and Ehop-016. All cells were infected equally whether or not they expressed ACE2 and all compounds significantly reduced SARS-CoV-2-pseudovirus mediated infection independent of ACE2 expression (a measured by % eGFP – expressing cells). At least 15,000 cells per condition were analyzed for n=3 replicates. *p<0.05, **p<0.01.

Gene set enrichment analysis (GSEA), using Kyoto Encyclopedia of Genes and Genomes (KEGG), of the mock vs PLOs 3 hpi revealed enrichment of pathways associated with Pyrimidine metabolism (*51*) and ECM receptor interaction (*52*) **[Fig. 2c]**, pathways previously associated with pulmonary infection (*53, 54*) while mock vs PLOs 24 hpi revealed enrichment pathways associated with oxidative phosphorylation (*55–57*) and steroid biosynthesis (*30*). Ingenuity pathway analysis (IPA) revealed associations with IL-17 signaling, pulmonary fibrosis idiopathic signaling, and IL-8 signaling **[Supplementary Fig. 4a].** Upstream regulators in these pathways aligned with signal transduction pathways controlled by IL-1, IFN, NF-B and TNF **[Supplementary Fig. 4b].**

These data emphasize that normal lung epithelial cells, without the intercession of hematopoietic-derived cells or a circulatory system, can initiate innate immune responses to SARS-CoV-2 infection, impact gene expression related to stress resistance, and influence the integrity and function of epithelial cells at this mucosal barrier (*58–60*).

To determine cell specific gene expression changes in the PLOs, we performed scRNA seq **[Fig. 2d, Supplementary Fig. 4c]** 16 hpi with the Alpha (hCoV-19/USA/CA_UCSD_5574/2020, B.1.1.7) variant. First, we identified infected cell clusters based on expression of viral gene transcripts. SARS-CoV-2 transcripts were detected in most clusters at various expression levels **[Fig. 2e]**; notably, cluster 14 had the highest expression of all the viral genes but was not representative of a specific cell type or function. Cluster 7 (ECM) had the next highest expression of the viral genes except for E, and clusters 9 (club cell) and 16 (alveolar-like cell) were the third highest, expressing approximately half of the viral genes. The genes of the structural proteins of SARS-CoV-2 (*61*), nucleocapsid (*N)* was expressed in all clusters, while membrane (*M*), envelope *(E)* and spike protein (*S)*, were more restricted. Accessory transcripts (*62*) *ORF10* showed expression in all clusters **[Fig. 2e]**. This finding is comparable to the temporal expression of individual SARS-CoV-2 genes after Calu3 cell infections (*63*).

To evaluate the correlation between viral gene expression and known viral entry genes, we next examined the gene expression of *ACE2* and *TMPRSS2* (*54, 64–72*) in conjunction with infected clusters **[Fig. 2f]**. Interestingly, ECM, alveolar-like cells, and the infected cluster showed the highest expression of the viral genes but the lowest expression of *ACE2* and *TMPRSS2*. We interrogated other viral entry genes (*73*) including *FURIN* (*74, 75*)*, HAVCR1* (*76*)*, AXL* (*77*)*, NRP1* (*78, 79*)*, SLC6A19* (*69, 80*)*, CTSB* (*81*)*, CTSL* (*82*)*, CTSS* (*83*), and *BSG* (CD147) (*84, 85*) **[Fig. 2f]**. We found *BSG* expressed in 16 of the 18 clusters and *CTSB* was expressed in 11 of the 18 clusters. *CTSB* and *CTSL* code for proteins important in *endocytic viral entry* (*81–83*). Serous and mucous-like cells expressed both the canonical and non-canonical viral entry genes (*ACE2, TMPRSS2, CTSB, CTSL*), while primordial lung progenitor, pre-goblet, club, and ionocyte expressed *BSG* only. Our gene expression findings of the tropism of SARS-CoV-2 were verified with immunofluorescence analysis. HBEC-ALI cells and dissociated lung organoids 24 hpi with pseudovirus-GFP showed cells co-expressing ACE2 receptor and the GFP signal from the pseudovirus-GFP, yet there were also some cells that only expressed GFP **[Fig. 2g]**. Dissociated LOs infected with the Delta and Omicron variants also showed similar tropism, with some cells co-expressing ACE2 and the viral NC, and others expressing only NC **[Fig. 2h]**. To further interrogate the non-canonical, non-ACE2 receptor-mediated endocytic route of SARS-CoV-2 into lung cells, we utilized compounds that block endocytosis (via micropinocytosis). Endocytosis is regulated by PI3K signaling (which plays a critical role in endosome/macropinosome closure and fission from the plasma membrane); therefore, we used Apilimod, a phosphatidylinositol-3-phosphate 5-kinase (PIKfyve) inhibitor (*86*) (*87*), the PI3K inhibitor LY294002 and Ehop-016 (an inhibitor of Rac1 that perturbs actin cytoskeleton dynamics). PIKfyve functions in distributing endocytic cargo (*88*). Apilimod blocked FITC-Dextran uptake into A549 and SK-LU-1 lung cells as efficiently as the PI3K inhibitor LY294002 **[Fig. 2i]**. ACE2 expression in A549 lung cells did not affect the extent of endocytosis **[Figs. 2j, 2k]** and did not affect the endocytic blocking effects of Apilimod, LY294002 and Ehop-016 **[Fig. 2l]**. Having validated Apilimod as a tool for discerning the presence of endocytosis, it was applied to A549 cells (with and without ACE2) to determine whether SARS-CoV-2 enters lung cells via that route. The lung cells were pretreated with the above-mentioned compounds and then infected with pseudovirus-GFP. All cells were infectable, whether or not they expressed ACE2, and all compounds significantly reduced the number of GFP-expressing cells, including Apilimod, independent of ACE2 expression **[Fig. 2m]**. The efficiency with which Apilimod blocked infection (comparable to the other endocytosis-blocking agents), would suggest that SARS-CoV-2 does enter pulmonary epithelial cells by non-canonical as well as canonical routes.

To compare the efficiency of reducing viral infection using endocytic blocking agents (Apilimod), cathepsin blocking agents (ONO5334) (*89*) and (as a positive control) agents that block viral replication (remdesivir) (*90*), we utilized a diverse set of 3D iPSC-PLOs and infected them with WA1 SARS-CoV-2. At 24 hpi, Apilimod reduced SARS-CoV-2 infection in PLOs derived from different donors to a degree comparable to remdesivir **[Supplementary Fig. 5a-b]**.

These data suggest that Apilimod may be a synergistic therapeutic option early in SARS-CoV-2 infection to block entry by the virus that attempts to circumvent ACE2/TMPRSS2-mediated routes. The failure of the cathepsin inhibitor ONO5334 (*91*) to block infection consistently suggests that the cathepsins may not be an ideal target for therapeutics.

### Acute infection with SARS-CoV-2 induces an autonomous intrapulmonary inflammatory cascade in bystander cells

To determine the relationship between infection and the intrinsic epithelial cell response to viral infection, we analyzed the scRNAseq data after infecting all the LO types with the WA1 SARS-CoV-2 variant **[Supplementary Figs. 6a-b]**. We scored the clusters based on the expression of the viral genes *N, M* and *ORF10.* Clusters 5 and 9 showed the highest infection score, while clusters further from 5 and 9 showed a lower infection score **[Fig. 3a]**. When looked at separately, the viral gene *N* (*92*) was highly expressed throughout all the clusters, followed by *M* (*93*)*, ORF8* (*94*), and *ORF7a* (*95*) **[Fig. 3b]**. We then assessed the expression of type I interferon-inducible genes using an open-access Interferome database generated from gene expression profiles of cells treated with type I or II interferon (*96, 97*). We generated an interferome score for each cluster, with significant elevations of interferome scores in all clusters, consistent with the presence of viral gene expression in all clusters **[Figs. 3b-d]**. Analysis of relative viral gene expression and interferome scores showed that absolute levels of viral transcript did not directly correlate with abundance of interferome gene transcripts. We then looked at specific examples of interferome gene expression profiles and found increased expression in *IFI27, IFI6, IFIH1, IFIT-1, -2, -3* and *IFITM1* **[Figs. 3e-f]**. Chemokines were also differentially expressed with most clusters showing increased expression except for the highly infected cluster 5 which constitutively expressed *CXCL1, CXCL2, CXCL3*, and *CCL20* at high levels **[Fig. 3g]**. The distribution of chemokine expression did not correlate with either interferome gene expression or viral gene expression. Finally, we performed a canonical pathway analysis, and interrogated the disease and biological functions and upstream regulators using IPA (*98*). The Coronavirus Pathogenesis pathway had positive activation z-scores in most of the clusters, while EIF2 signaling had negative activation z-scores (*99*) **[Fig. 3h]**. Genes important in apoptosis and necrosis were increased (*100*) **[Fig. 3i]**, as were the upstream regulators of these canonical pathways *IFNL1, IFNA2, NONO, IRF7* and Interferon alpha (*101*) **[Fig. 3j]**. The correlation of viral gene expression with these inflammatory mediators was also conducted, and again showed non-overlapping profiles.

**FIGURE 3:**
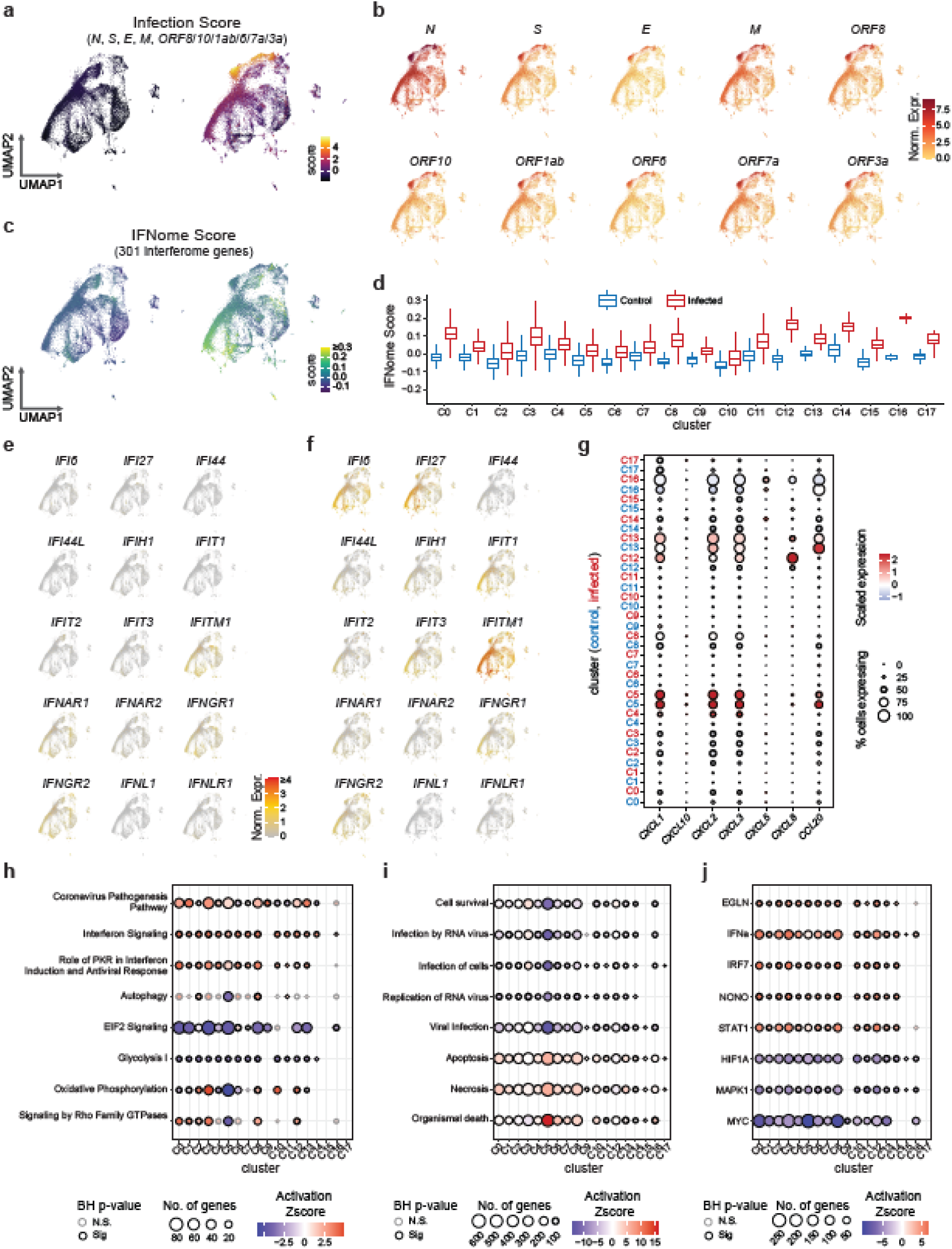
Acute infection with SARS-CoV-2 induces an autonomous intrapulmonary inflammatory cascade in bystander cells. **(a)** UMAP of infection score showing the highest expression of SARS-CoV-2 genes in clusters 5 and 9. The significance of those clusters is described in the text. Gradient shows highest expression levels in red. **(b)** UMAP of viral gene expression. All viral genes expressed in clusters 5 and 9, while viral gene *N* (nucleocapsid) is expressed in all clusters. Gradient shows highest expression levels in red. **(c)** UMAP depicting Interferome-I (IFN-I) score for each cluster. Highest expression is seen in clusters 0, and 12-17. Clusters with the highest IFN-I scores did not overlap clusters with the highest infection scores, indicating that non-infected “bystander” lung cells were being induced by infection of their lung cell “neighbors” to mount an IFN response. (d) Violin plots depicting a higher IFN-I score for each cluster when comparing mock vs. infected lung organoids. (e, f) UMAPs of a subset of interferon genes differentially expressed in (e) mock vs (f) infected LOs. Gradient shows highest expression levels in red. **(g)** Dot plot showing differentially expressed chemokine and cytokine genes by cluster comparing infected vs mock LOs. See text for a description of the significance of each. **(h)** Canonical pathway analysis of infected vs mock LOs by cluster using Ingenuity Pathway Analysis (IPA). Interferon signaling, Role of hypercytokinemia/hyperchemokinemia and Coronavirus Pathogenesis pathways had positive z-scores, while glycolysis, EIF2 signaling and gluconeogenesis had mostly negative z-scores. Oxidative phosphorylation was mixed. **(i)** Disease and Biological Functions using IPA. Viral infection, Replication of Virus and Infection of Cells had negative z-scores, while Organismal death and Necrosis had positive z-scores. These scores might be interpreted as the lung’s intrinsic/autonomous inflammatory viral defense system successfully impairing the continuation of the viral life cycle within the lung post-infection. **(j)** Upstream Regulators using IPA. *IFNL1, IFNA2, NONO, IRF7* and Interferon alpha had positive z-scores (consistent with the presence of an intrinsic intrapulmonary inflammatory feedback system) while *MYC, HIF1A* and *MAPK1* had negative z-scores.

Taken together, these data suggest that specialized lung cells are equipped to respond themselves to infection with different innate immune mechanisms. These data further suggest a threshold for viral expression that triggers differential expression of the interferome and chemokines in human lung cells without the intercession, or even presence, of hematopoietic derivatives or a vascular compartment.

### Natural surfactant and recombinant SP-B reduce SARS-CoV-2 infection and cell death

Having demonstrated the broad susceptibility of specialized human lung cells to SARS-CoV-2, and their unique ability to promote innate immune responses and recruit inflammatory cells by chemokine production, we returned to the role of surfactants during infection. To understand the effect of surfactant on the susceptibility of a cell to infection by SARS-CoV-2, we first used immunofluorescence to probe for the four surfactant proteins (SPs) 24 hpi in the WLOs and DLOs; we detected co-expression of viral NC with SP-A**+**, SP-B**+**, SP-C**+**, and SP-D**+ [Fig. 4a]**. Intriguingly, only infected cells expressed SP-C and SP-D while both infected and uninfected bystander cells expressed SP-A and SP-B. But, of the SPs, SP-B drew our greatest attention, because its presence (complexed with the lipid DPPC), is the *sine qua non* for pulmonary surfactant function, the substance required for ventilation and gas exchange in the alveoli. SP-B is required for normal lamellar body biogenesis, which is the storage form of surfactant (*102–104*). It also regulates SP-C post-translational modification (*105*). As noted previously, if preterm babies are born prior to the onset of SP-B synthesis, neonatal RDS develops. If SP-B fails to be produced at all, as in rare cases of mutations in the SP-B gene, lethal respiratory failure ensues even in full-term infants, reversible only by lung transplantation (*106*).

**FIGURE 4:**
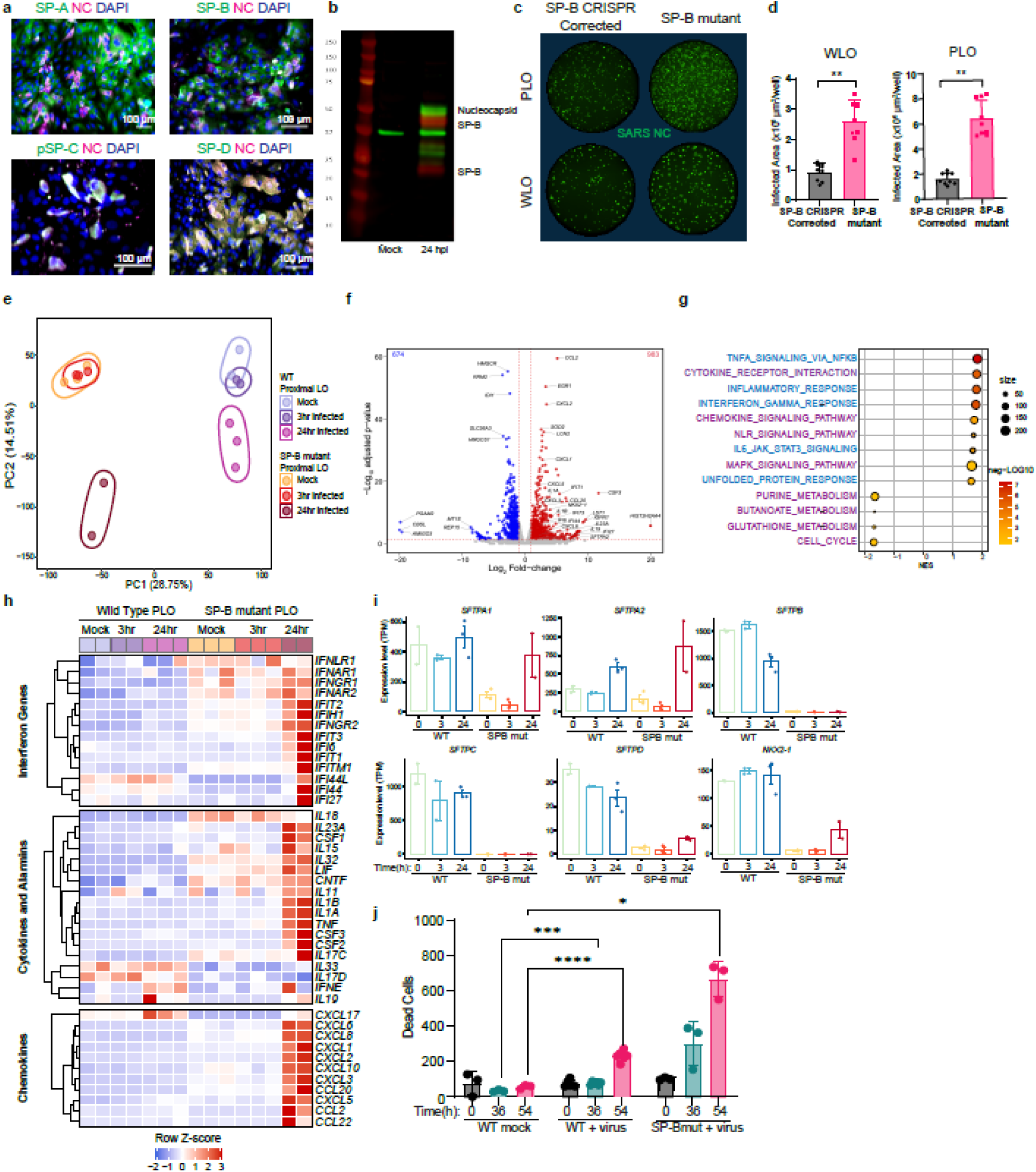
Surfactant suppresses viral infection and modulates inflammation. **(a)** Immunostaining of dissociated 3D PLOs (acutely dissociated for better visualization of individual cells), 24 hpi, showing the expression of surfactant proteins (SPs) A, B, C and D in many lung cell types. **(b)** Western blot of Mock and 3D PLOs, 24 hpi. Infected organoids show expression of the SARS-CoV-2 nucleocapsid (NC) protein as well as 2 isoforms of SP-B including pro-form (25 kD) and pre-pro form (42 kD). **(c)** Immunostaining of the SARS-CoV-2 NC in a 96 well format of LOs from an SP-B deficient patient (due to a frameshift mutation) and the same LOs corrected via CRISPR genomic editing to enable SP-B production. Infection was performed at an MOI of 0.1 and the wells were overlayed with carboxymethylose to slow down viral spread over 24 hours. Note that the absence of SP-B expression enabled much greater SARS-CoV02 infection of the same isogenic LOs than when the SP-B gene expression was restored. Experiment was repeated 9 times and **(d)** graphed as total infected cell area. **** = p-value < 0.0001. **(e)** PCA plot of WT PLOs mock infection (lavendar), 3 hpi (purple), 24 hpi (pink) and SP-B mutant PLOs mock infection (orange), 3 hpi (red) and 24 hpi (brown). Note that infected LOs lacking SP-B segregate dramatically from those expressing SP-B and that a longer post-infection period (24 hours) in the absence of SP-B creates another distinct cluster (see text for details and explanation). **(f)** Volcano plot analysis of differential gene expression of SARS-CoV-2 infected SP-B deficient PLOs versus mock infection. The red lines indicate a p-adjusted value <0.05. The absence of SP-B enables an upregulation of numerous toxic cytokines (see text for details). **(g)** Gene over-representation analysis using Kyoto Encyclopedia of Genes and Genomes (KEGG) pathway database of SARS-CoV-2 24 hpi vs mock infected SP-B mutant PLOs. **(h)** Heat map showing average expression (represented as z-score) of key innate and inflammatory genes in wild type and SP-B deficient 3D PLOs at 0, 3 and 24 hpi. Inflammatory pathways are heavily upregulated in the latter compared to the former (see text for details). The individual numerical value per condition is listed in Table 1. **(i)** Surfactant proteins and key transcription factor *NKX2-1* expression in wild type (WT) and SP-B deficient 3D PLOs at 0, 3 and 24 hpi. Note that the expression of *SFTPA1* and *SFTPA2* increase in both WT and SP-B deficient PLOs, 24 hpi. The data are representative of three independent experiments. **(j)** SP-B deficiency impairs cell viability within LO cells following SARS-CoV-2 infection compared to WT LOs. Both types of LOs were infected with SARS-CoV-2 at MOI = 0.1 and propidium iodide (PI) (a marker of cell death) expression was tracked over 54 hours. Significance calculated using 2-way ANOVA. * = p-value < 0.05, *** = p-value < 0.001, **** = p-value < 0.0001. SP-A = Surfactant Protein A; SP-B = Surfactant Protein B; SP-C = Surfactant Protein C; SP-D = Surfactant Protein D; NC = Nucleocapsid; WLO = Whole Lung Organoid; PLO = Proximal Lung Organoid; MUT = Mutant

As noted above, AT2 cells in the LOs produced SP-B normally, as do secretory cells (club cells) in the airways. On western blots of PLOs 24 hpi, we detected 2 isoforms of SP-B in these cells: 42kDa and 25kDa (*11*) **[Fig. 4b]**. AT2 cells produced SP-B at 8kDa (as they would in vivo for surface tension reduction) (*107*). The induction of SP-B post-viral infection in pulmonary cells, particularly those that normally do not produce surfactants *in vivo* for facilitating ventilation, suggests that SP-B may be induced by pathogenic viruses for anti-viral, homeostatic and/or anti-inflammatory purposes (*108–110*).

To explore the specific roles of SP-B, we not only included the above-mentioned normal hiPSC-derived LOs, but also LOs generated from hiPSCs from a patient with a lethal mutation in *SFTPB p.Pro133GlnfsTer95 (PRO133)* (*111, 112*). This mutation is a homozygous, loss-of-function mutation that results in a frameshift leading to an early stop codon. The transcript is unstable and thus does not translate a protein product (*20*). We also obtained LOs from these same hiPSCs (hence, isogenic) in which this mutation was corrected via CRISPR-mediated genome editing (*113*). The LOs were acutely dissociated and infected with the Beta variant (hCoV-19/South Africa/KRISP-K005325/2020, B.1.351, Beta) of SARS-CoV-2 at an MOI of 0.1 as a monolayer to provide adequate and homogeneous access of all cells to the virus. At 24 hpi, the cells were fixed, and NC immunoreactivity quantified **[Fig. 4c]**. SP-B-mutant cells had higher numbers of infected cells compared to isogenic corrected cells **[Fig. 4d]**. These data suggested that the absence of SP-B impairs resistance to SARS-CoV-2 infection, possibly by providing a physical barrier to viral infection in the lung.

To investigate how SP-B may be impacting viral entry and subsequent dissemination to neighboring cells, we examined SP-B-mutant PLOs compared to normal PLOs by bulk RNA-sequencing at 0, 3, and 24 hpi with the Delta variant of SARS-CoV-2. Strikingly, PCA demonstrated that SP-B-mutant PLOs at 24 hpi occupied a distinct transcriptional space from all the wild type (WT) samples and the mock and 3 hpi SP-B mutant samples **[Fig. 4e]**. We determined the proportion of cells infected by examining the viral reads and found that 16-20% of the SP-B mutant PLOs 24 hpi were infected compared with less than 1% of normal PLOs, 24 hpi **[Table 1]**.

**Table 1.** SARS-CoV-2 viral reads from uninfected and infected proximal lung organoids.

| **Group** | **Viral Reads (%)** |
| --- | --- |
| SP-B Mutant PLO uninfected | 0 |
| SP-B Mutant PLO uninfected | 0 |
| SP-B Mutant PLO uninfected | 0 |
| SP-B Mutant PLO + Delta variant 3 hpi | 0.11 |
| SP-B Mutant PLO + Delta variant 3 hpi | 0.38 |
| SP-B Mutant PLO + Delta variant 3 hpi | 0.40 |
| SP-B Mutant PLO + Delta variant 24 hpi | 20.41 |
| SP-B Mutant PLO + Delta variant 24 hpi | 16.48 |
| Wild type PLO uninfected | 0 |
| Wild type PLO uninfected | 0 |
| Wild type PLO uninfected | 0 |
| Wild type PLO + Delta variant 3 hpi | 0.14 |
| Wild type PLO + Delta variant 3 hpi | 0.36 |
| Wild type PLO + Delta variant 24 hpi | 0.68 |
| Wild type PLO + Delta variant 24 hpi | 0.25 |
| Wild type PLO + Delta variant 24 hpi | 0.58 |

PLO = Proximal Lung organoid; hpi = hours post infection

Infected SP-B-mutant PLOs were characterized by increased chemokine transcripts (*CCL20, CXCL-1, -2, -3, -5* and *-6*), interleukin transcripts (*IL1A, IL1B, IL19, IL23A*), interferon-inducible genes, and other cytokines including *CSF3***[Fig. 4f and Supplementary Figs. 7a,b]**. The surfactant associated genes *SFTPA2,* and *NKX2-1* were also significantly induced, in contrast to the above-mentioned normal PLOs. GSEA, (using KEGG), of the mock vs. 24 hpi SP-B-mutant PLOs were characterized by activation of pathways associated with Cytokine Receptors, Chemokine Signaling, NOD-like receptor signaling, Interferon signaling, and MAPK Signaling **[Fig. 4g]**. IPA was consistent with the WT PLO data except that the average expression of the canonical pathways **[Supplementary Fig. 4a]** and upstream regulators **[Supplementary Fig. 4b]** was greater in the SP-B mutant PLOs. Even the IFNome 90 enrichment scores were higher in the SP-B mutant PLOs compared to the WT PLOs **[Supplementary Fig. 4d]**.

Although the uninfected SP-B-mutant PLOs had increased expression of some interferon-inducible genes, we found significant differences in the basal levels of the key *protective* interferon-inducible genes *IFI44L* and *IF127,* which were higher in the normal PLOs compared to the SP-B mutant PLOs **[Fig. 4h]**. This finding reinforced our speculation that SP-B played a homeostatic role in maintaining lung function by modulating inflammatory gene expression at steady state. Furthermore, cytokines *IL33* (*114*)*, IL17D*(*115*), and *IL19* (*116*) were expressed at baseline in normal PLOs but were not expressed in the SP-B-mutant PLOs. Also, mediators that correlate with *severity* of SARS-CoV-2 infection - *IL1A, IL1B, CCL2, CXCL8, CXCL10*, and *CSF3* - were induced in SP-B mutant PLOs 24 hpi (*117*).

To investigate further the role of SP-B in gene regulation, we measured the expression of SP genes and their transcription factor *NKX2-1* (*118, 119*). The normal PLOs had higher expression of all the surfactant protein genes and *NKX2-1* pre- and post-infection (0, 3, and 24 hpi) compared to the SP-B-mutant PLOs (except for *SFTPA2*, which had the highest expression in 24 hpi SP-B-mutant PLOs) **[Fig. 4i]**.

To determine viral tropism in the SP-B mutant PLOs, we performed scRNAseq **[Supplementary Figs. 4c and 7]** 16 hpi with the Alpha variant. The cell cluster types were comparable to the WT PLOs **[Supplementary Fig. 7a]**. SARS-CoV-2 transcripts were detected in most clusters at various expression levels, notably cluster 14, which had the highest expression of all the viral genes but was not representative of a specific cell type or function, consistent with findings in the infected WT PLOs **[Supplementary Fig. 7b]**. Interestingly, the viral gene *N* was expressed in all clusters in the WT PLOs **[Fig. 2e]**, but, in the SP-B mutant data, the pre-goblet cells and the serous cells did not express *N*. In fact, the pre-goblet cells did not express any viral genes. The ECM cells did not highly express the viral genes in the SP-B mutant data set, but the alveolar-like cells had even more viral transcripts compared to the WT PLOs. No viral genes were expressed in all the clusters in the SP-B mutant PLOs. *N* was expressed in 16 out of 18 clusters, and *ORF10* was limited to 10 out of 18 **[Supplementary Fig. 7b]**. The viral entry receptors also differed between the WT and SP-B mutant data. ACE2 and TMPRSS2 were limited to 1-2 clusters in the infected WT PLOs **[Fig. 2f]**, but, in the SP-B mutant data, they were highly expressed in lung progenitors, alveolar type 2 cells, club cells, and ionocytes. The other viral entry genes were consistent between the WT and SP-B mutant PLOs except that club cells expressed more transcripts of *CTSS, CTSL*, and *CTSB* **[Supplementary Fig. 7c]**.

Taken together, these data suggest that the absence of SP-B in human pulmonary epithelial cells abets greater viral infection, suppresses baseline anti-viral defense mechanisms, and enables a more inimical post-viral inflammatory response.

### Viability and viral dissemination in LOs following SARS-CoV-2 infection is reduced by BH3 mimetics and surfactant protein B

To evaluate if altered virus gene expression in SP-B mutant PLOs also impacts cell viability, we tracked cell death over 90 hours and found that the SP-B mutant PLOs showed higher rates of cell death after 36 hours, compared to WT PLOs **[Fig. 4j]**. These data reflect the increased viral gene expression in SP-B-mutant LOs, but also reveal an unexpected role for SP-B in viral host defense and maintenance of lung cell viability and function during acute stages of viral infection prior to recruitment of hematopoietic cells into the lung. However, to better understand SP-B’s mechanism-of-action in this viral pathogenic process, we needed to better understand the type of cell death that SP-B was ameliorating.

We have previously shown that SARS-CoV-2 infection-induced endoplasmic reticulum (ER) stress and an unfolded protein response (UPR) triggers caspase-mediated apoptosis (*120*). This process was evident in post-mortem human lung tissue from patients with COVID-19, and in SARS-CoV-2-infected Vero E6 cells – an African Green Monkey kidney epithelial cell line deficient in interferon production (*121*). The kinetics of cell death triggered by SARS-CoV-2 could be modulated by BH3 mimetics and inhibited by caspase antagonists, consistent with the role of Bcl-2 family proteins as regulators of cell viability and apoptosis during SARS-CoV-2 infection (*120*). To understand the role of virus-triggered apoptosis in human lung cells, we infected hiPSC-derived PLOs and then tracked the changes in viability in cells treated with caspase inhibitors or BH3 mimetics. To enable tracking of the rapid changes in viability induced by SARS-CoV-2 infection, 3D hiPS-derived PLOs were acutely dissociated and plated as a monolayer and then infected with SARS-CoV-2. Virus infection resulted in death of 3D organoids, as well as freshly dissociated organoids infected as a monolayer of cells **[Fig. 5a]**. Caspase inhibition with Q-VD-OPh blocked SARS-CoV-2-induced death of human lung cells, consistent with a role for apoptotic caspases in cell death of human lung cells. Addition of BH3 mimetics to cultures of SARS-CoV-2-infected human lung cells accelerated cell death, again consistent with a role for Bcl-2 family proteins in regulating cell viability **[Figs. 5b, c]**. Inhibition of Bcl-2, Bcl-x_L_, and Bcl-w with ABT-737 increased cell death of infected cells, whereas the selective BH3 mimetics ABT-199, an antagonist of Bcl-2, and S63845, an antagonist of Mcl-1, caused only minor reductions in viability compared to infected controls. We then tested A-1331852, a selective and potent antagonist of Bcl-x_L_, to understand its contribution to the changes in viability seen with ABT-737. Inhibition of Bcl-x_L_ accelerated the death of SARS-CoV-2-infected human lung cells **[Fig. 5d]**. These data confirmed a vital role for Bcl-2 family proteins, and in particular Bcl-x_L_, in control of human lung cell viability and demonstrated synergistic impacts on cell viability in the presence of SARS-CoV-2.

**Figure 5.**
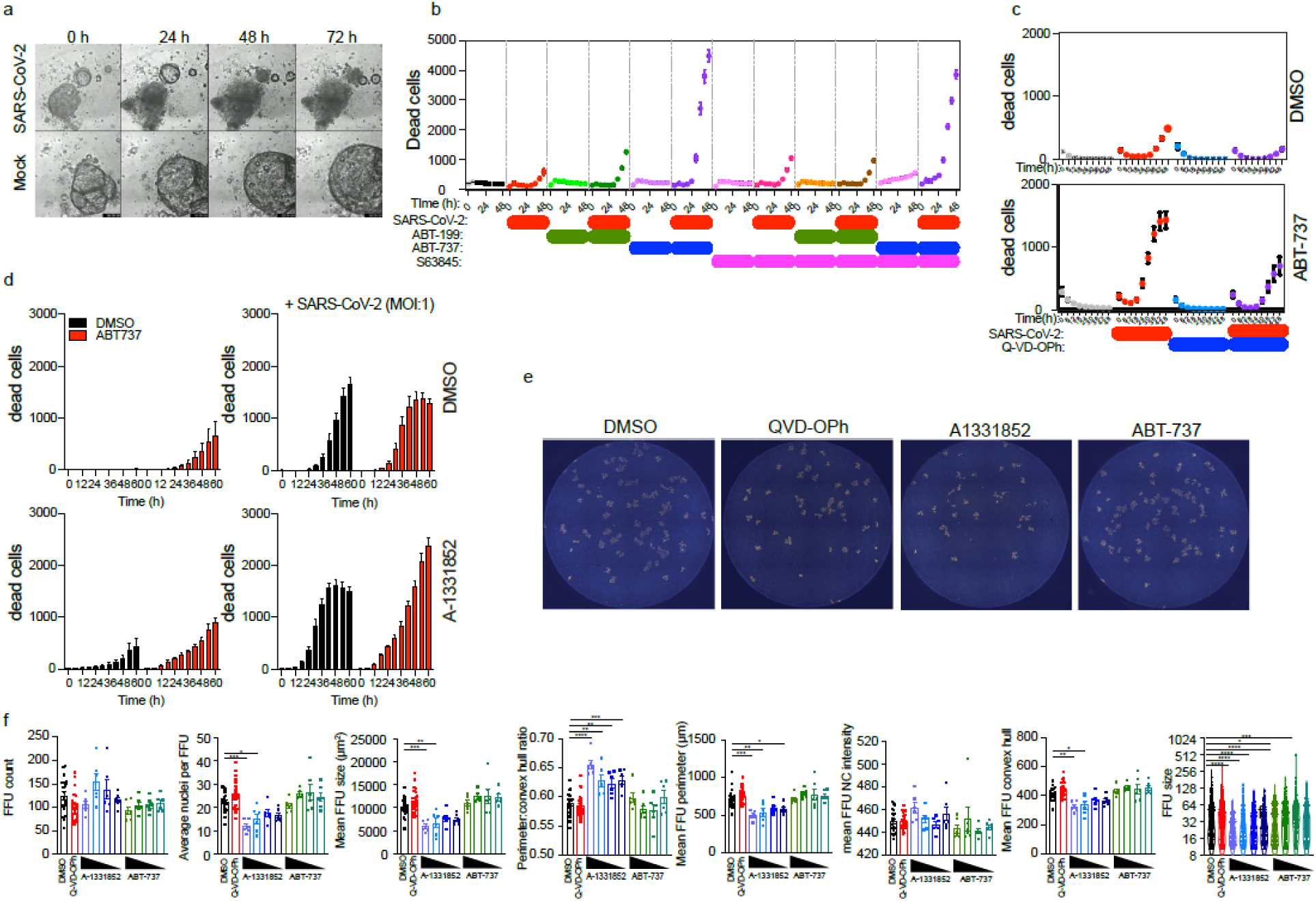
Viability and viral dissemination in lung organoids (LOs) following SARS-CoV-2 infection is reduced by BH3 mimetics and caspase antagonists. **(a)** Live cell imaging of 3D LOs by phase contrast imaging showing morphological changes triggered by SARS-CoV-2 (MOI 1) infection or mock infection controls. Images collected every 24h by Incucyte S3. Note progressive dissolution of integrity of the LOs. **(b-c)** SARS-CoV-2 infection of human lung cells triggers a reduction in cell viability. Viability of monolayers of dissociated 3D LOs was tracked using propidium iodide (PI). Cells were infected with SARS-CoV-2 (MOI:0.25) for 1h and then treated with 125 nM ABT-737, ABT-199, S-63845, or Q-VD-OPh (10uM). Viability was monitored every 6h using an Incucyte S3. Data representative of 3 independent experiments. Note that caspase inhibition with Q-VD-OPh blocked SARS-CoV-2-induced death of human lung cells, consistent with a role for apoptotic caspases in cell death of human lung cells. Addition of BH3 mimetics to cultures of SARS-CoV-2-infected human lung cells accelerated cell death, consistent with a role for Bcl-2 family proteins in regulating cell viability and apoptosis during SARS-CoV-2 infection. Inhibition of Bcl-2, Bcl-xL, and Bcl-w with ABT-737 increased cell death of infected cells, whereas the selective BH3 mimetics ABT-199, an antagonist of Bcl-2, and S63845, an antagonist of Mcl-1, caused only minor reductions in viability compared to infected controls. **(d)** The Bcl-xL-selective inhibitor A-1331852 synergized with SARS-CoV-2 (MOI:0.25) to reduce cell viability of human LO cells, confirming a vital role for Bcl-2 family proteins, and in particular Bcl-xL, in control of human lung cell viability, particularly in the face of SARS-CoV-2 infection. **(e)** Quantitative fluorescent focus unit (FFU) immunoassay (developed to record the size of foci and the number of infected cells within each focus) of SARS-CoV-2 infected VeroE6 cells (MOI:0.01) exposed to ABT-737 (BCL-2 inhibitor) (250nM-2uM), A-1331852 (Bcl-xL antagonist) (250nM-2uM), or Q-VD-OPh (caspase inhibitor) (10uM), and overlayed with 1.2% carboxymethylcellulose in media to inhibit viral spread. Immunostaining of SARS-CoV-2 nucleocapsid (NC) in 96 well plates and counterstained with Hoechst. Representative of 3 independent experiments. **(f)** Quantitative image analysis of individual foci indicating the focus count in individual wells, the average number of infected cells per focus in each well, and the average size, perimeter, and convex hull of foci in each well. Focus size reflects the number of infected cells arising from dissemination of virus from individual infected cells within a focus. n=3. *p < 0.05, ***p < 0.001,****p < 0.001. Kruskal-Wallis test. Note that treatment of virus-infected cultures with A-1331852 resulted in a reduction in size of foci of infection as assessed by the number of infected nuclei per cluster, the FFU cluster area, the FFU cluster perimeter, and the FFU cluster convex hull. Inhibition of Bcl-xL did not change the number of foci or viral NC levels within the FFU cluster as assessed by the average fluorescence intensity of intracellular NC. Taken together these data suggest that rapid induction of apoptosis of virus-infected human lung cells occurs at an early phase of the viral life cycle and that apoptosis induces lung cells inherently to interfere with further virus replication and dissemination, an intrinsic self-defense mechanism.

The death of a virus-infected cell serves as an endpoint for virus replication within that cell, but it is not known if apoptosis prevents viral dissemination to surrounding cells. Virus infection results in transcriptional changes in cell death regulators of human lung cells, including Bcl-2 family proteins, suggesting that regulated cell death may serve to restrict viral dissemination. We hypothesized that rapid induction of apoptosis using BH3 mimetics would restrict virus dissemination by selectively targeting infected cells primed for apoptotic cell death. We developed a quantitative fluorescent focus unit (FFU) assay to record the size of foci and the number of infected cells within each focus. Using a custom-scripted Image J code for automated image analysis, and a custom R Shiny tool called FFUTrackR for data visualization of the analysis output, we assessed characteristics of fluorescent foci in virus-infected cultures **[Fig. 1d]**.

Treatment of virus-infected cultures with the Bcl-x_L_ antagonist A-1331852 resulted in a reduction in size of foci of infection as assessed by the number of infected nuclei per cluster, the FFU cluster area, the FFU cluster perimeter, and the FFU cluster convex hull. Inhibition of Bcl-x_L_ did not change the number of foci or viral NC levels within the FFU cluster as assessed by the average fluorescence intensity of intracellular SARS-CoV-2 nucleocapsid **[Figs. 5e-f]**. These data suggest that rapid induction of apoptosis of virus-infected cells can occur at an early phase of the viral life cycle that interferes with virus replication and dissemination. In contrast, treatment with Q-VD-OPh did not alter virus dissemination suggesting that apoptotic caspases do not inactivate virus, nor are they required for virus dissemination **[Figs. 5e-f]**. These data also indicated that induction of apoptosis which is slower than that induced by A-1331852 - including that by ABT-737, ABT-199, and S63845 - fails to prevent virus dissemination, and likely releases pre-formed virions. Indeed, the spread of virus from dying cells to neighboring cells is the basis for the plaque assay. These data indicate that apoptosis can restrict viral dissemination, but only if it is engaged before significant viral replication occurs in the cell. In other words, taken together these data suggest that rapid induction of apoptosis within infected lung cells at an early phase of the viral life cycle induces the lung cells inherently to interfere with further virus replication and dissemination, an intrinsic self-defense mechanism.

Furthermore, based on this understanding of SARS-CoV-2-mediated cell death, we can impute that SP-B, based on its restriction of viral spread, may also prevent widespread apoptotic death of infected lung cells to preserve lung function in the acute phase of SARS-CoV-2 infection prior to significant hematopoietic cell recruitment.

### SP-B mutant gene correction, recombinant SP-B, and exogenous surfactant all reduce SARS-CoV-2 infection

We have shown that LOs lacking SP-B, the *sine qua non* of functional surfactant, have increased rates of SARS-CoV-2 infection and apoptotic cell death, presumably because surfactant not only provides a barrier to infection but mediates an inflammatory host defense that combats virus survival and dissemination. This speculation was supported by our observation that CRISPR-mediated correction of a deletion mutation of the gene encoding SP-B reversed these processes. Furthermore, adding recombinant SP-B (rSP-B) to SP-B-mutant LOs similarly reduced infection and cell death: when SP-B-mutant LOs were incubated in rSP-B (5µM) prior to infection, cell death (at 36 and 54 hpi) **[Fig. 6j]** was reduced.

**FIGURE 6:**
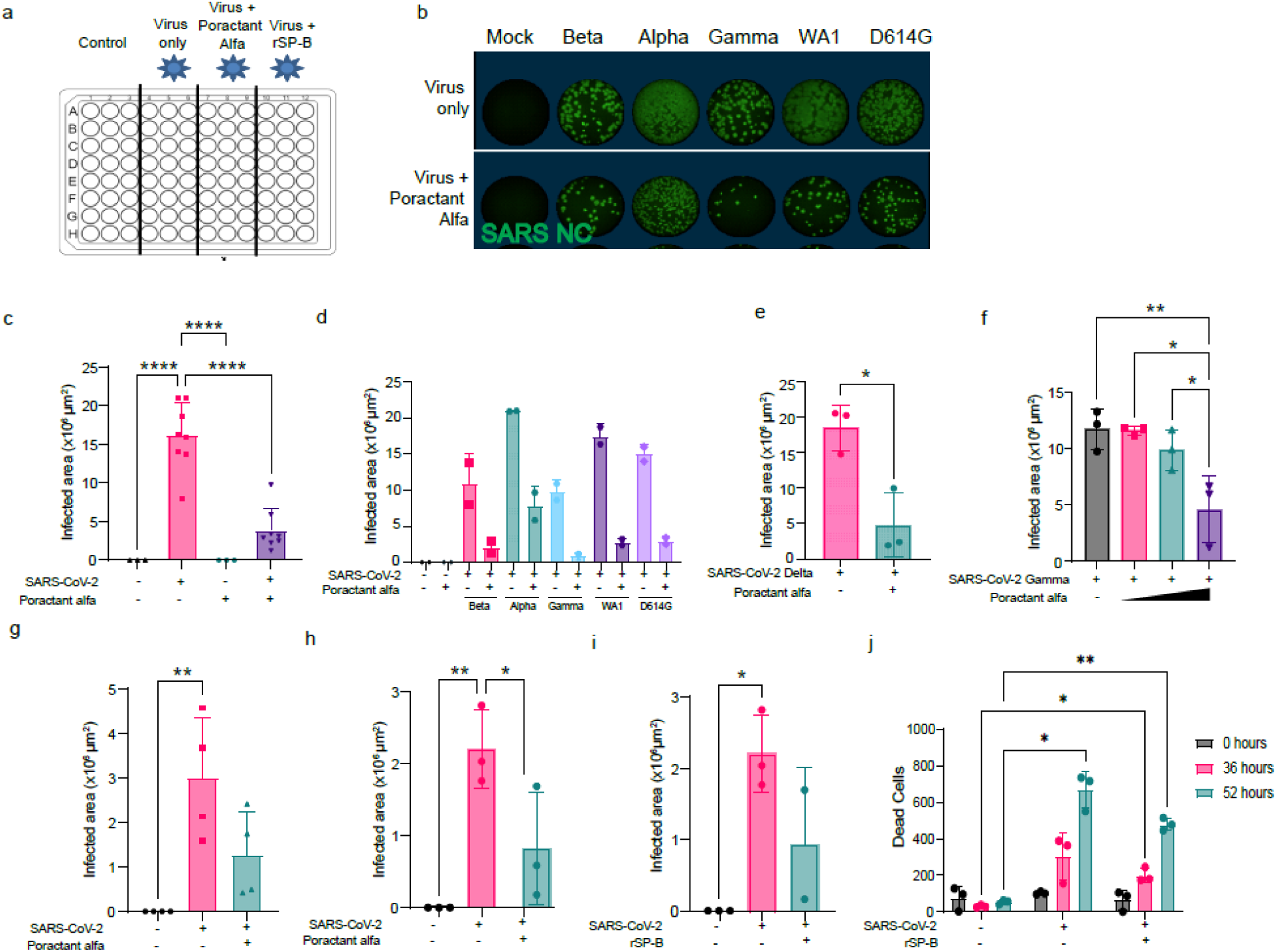
Natural surfactant and recombinant SP-B reduce SARS-CoV-2 infection and cell death. **(a)** Schematic of infection of dissociated lung organoids (LOs) or Vero E6 cells treated with FDA-approved, clinically-used, commercially-available porcine-derived surfactant, “poractant alfa” (1mg/ml) or recombinant surfactant protein B (rSP-B, 5µM). Vero E6 cells or LO derived cells were exposed to the surfactants for 30 minutes prior to infection. SARS-CoV-2 at an MOI of 0.1 was added to each 96 well plate (excluding control cells) for one hour. Cells were then overlayed in media supplemented with 1.2% carboxymethylcellulose and infected for 24-72 hours. **(b)** Immunofluorescent whole well scan of Vero E6 cells 24 hpi with different variants of SARS-CoV-2 with or without the addition of poractant alfa. Wells were stained for the nucleocapsid (NC) protein intracellularly to quantify degree of viral infection and quantified (FFU). **(c)** Quantification of infected Vero E6 cells 24 hpi (all variants taken together) with or without the addition of poractant alfa. Note that surfactant dramatically diminishes the extent of viral infection. **(d, e)** Quantification of degree of infection of Vero E6 cells (24 hpi) based on specific SARS-CoV-2 variant, with or without the addition of poractant alfa; (d) shows the results from variants Beta, Alpha, Gamma, WA1, and D614G; (e) shows the results from the Delta variant. **(f)** Total infected Vero E6 cells 24 hpi with the SARS-CoV-2 Gamma variant after different dosages of poractant alfa, showing its antiviral actions to increase with dose escalation. **(g-j)** The addition of exogenous surfactant or recombinant SP-B (rSP-B) to SP-B deficient LOs can rescue those LOs from their otherwise higher susceptibility to viral infection and cell death (see Fig. 4). **(g)** Reiteration of the normal inhibition of infection of WT PLO cells by the SARS-CoV-2 Gamma variant with the addition of poractant alfa. Total infected SP-B-deficient distal LO cells, 24 hpi with the Gamma variant, is diminished by the addition of poractant alfa **(h)** or rSP-B **(i).** Similarly, cell death (quantified using PI) of infected SP-B-deficient PLOs is reduced at 52 hours by the addition of rSP-B **(j).** Significance calculated using 2-way ANOVA * = p-value < 0.05, ** = p-value < 0.01. **** = p-value < 0.0001

We next explored directly clinically-relevant implications of these observations. We tested whether an FDA-approved, naturally-derived, exogenous whole surfactant preparation which is presently in routine clinical practice, could also inhibit or reverse the pathogenic actions of SARS-CoV-2 **[Fig. 6a]**. First, we performed dose-finding and timing studies by applying the commercially available porcine-derived surfactant, poractant alfa (*122*) **[Supplementary Table 2]** to Vero E6 cells 30 minutes prior to infection with the SARS-CoV-2 variants WA1, Alpha, Beta, Gamma (hCoV-19/Japan/TY7-503/2021, P.1), and D614G (USA/NY-PV08410/2020, B.1) (**[Fig. 6b]**. Based on assessments of NC immunopositivity 24 hpi **[Supplementary Fig. 6]**, there was a significant reduction in the number of SARS-CoV-2-infected cells in surfactant-treated cells compared to untreated **[Fig. 6d, e]**. We observed a reduction in the number of infected cells with increasing concentrations of surfactant **[Fig. 6f]**. We then pre-treated infected DLOs with poractant alfa and similarly found a significant reduction in the number of infected cells **[Fig. 6g, h]**. To confirm that the underlying mechanism of therapeutic action of this practical clinical intervention pivoted on the expression of SP-B, we confirmed that adding recombinant SP-B (rSP-B) alone to SP-B-mutant LOs replicated this reduction in infection: when SP-B-mutant LOs were incubated in rSP-B (5µM) prior to infection, SARS-CoV-2 infectivity (24 hpi) was equally suppressed **[Fig. 6i].**

## DISCUSSION

Here, we employed an experimental system capable of determining – prospectively and in an unbiased manner – the tropism of SARS-CoV-2 in the lung, mapping at a single cell level the downstream transcriptional, proteomic, and functional response of pulmonary epithelial and mesenchymal cells upon initial viral exposure. In addition, this system can begin to address concerns regarding COVID-19 disparity because the lung organoids (LOs) have been derived from men and women from a range of racial groups.

In employing this system, we observed multiple unexpected responses of the lung to SARS-CoV-2 infection – with implications for the lung’s response to viral infection more generally. We noted that there exists – as part of the lung’s inherent anti-viral host defense mechanism – an under-appreciated intrinsic, autonomous intrapulmonary post-infection inflammatory cascade that is mediated, at least in part, by surfactant protein B (SP-B). In other words, it appears that surfactant plays a role beyond its well-studied physical-chemical one of reducing alveolar surface tension to maximize ventilation and oxygenation. SP-B provides a pivotal barrier to viral entry and persistence and regulates virus-induced intrapulmonary inflammation-related gene expression which further mitigates pathogen survival and dissemination.

In support of this conjecture was, first, our surprising observation that the intracellular expression of SARS-CoV-2 genes and proteins (signs of viral entry and replication) was greatest in SP-B mutant LOs. It bears reiterating that, because our LOs are derived from patient-specific hiPSCs, we were able to construct LOs from a patient afflicted with a lethal deletion mutation in the SP-B gene p.Pro133GlnfsTer95 (PRO133) precluding all surfactant protein production. As a control (in addition to WT hiPSC derived LOs), we used LOs from the same patient with SP-B function and surfactant production restored via CRISPR-Cas9-mediated correction of the mutation. As additional complementation controls in some experiments, we administered recombinant SP-B or exogenous complete surfactant. Dexamethasone is a strong anti-inflammatory drug that also induces AT2 cell maturation and surfactant production **[Fig. 1A, Supplemental Fig. 1a, Supplementary Movie 1, Methods]**. Dexamethasone has shown benefits clinically in reducing morbidity and mortality in patients with CARDS, its efficacy typically attributed to its anti-inflammatory and anti-hypotensive actions; its potential actions in reducing viral entry and dissemination in pulmonary epithelial cells via its impact on AT2 cell regeneration and/or surfactant production may also bear further investigation (beyond the scope of this study).

The expression of the surfactant protein genes was dynamic in the WT LOs. *SFTP-A, C* and *D* initially decreased 3 hpi, and then increased 24 hpi. Only *SFTPB* increased at 3 hpi then expression was reduced at 24 hpi, indicating a higher level of transcription earlier in the infection. The SP-B mutant LOs had minimal expression of the surfactant protein genes, with only *SFTPA* increasing significantly at 24 hpi. *SFTPB* is controlled by multiple upstream regulators, including inflammatory mediators IL1, IL6, TGFβ and the transcription factors *NKX2-1, JAK1* and *STAT3* **[Supplementary Fig. 7]**. Therefore, absence of SP-B may exacerbate the deleterious components of the post-infection inflammatory response leading to increased pulmonary cell death and apoptosis. What makes this observation striking is the fact that the LOs contained no cells of hematopoietic or mesodermal origin, i.e., no alveolar or systemic immune cells. Therefore, these data begin to suggest that, in response to SARS-CoV-2 infection, lung epithelial cells *themselves* might serve as initiators of innate immunity and impact gene expression related to oxidative stress tolerance and barrier cell maintenance. Broadly stated, under normal circumstances (as seen in WT LOs with normal SP-B levels), constant low-level intrapulmonary immune surveillance is operative even pre-infection, as reflected in baseline detectable levels of IL-33, IL-17, and expression of receptors for type I, II, and III interferons **[Fig. 5h]**. However, in SP-B mutant LOs, those surveillance levels are abnormally low. Upon infection with SARS-CoV-2, while in WT LOs those inflammatory response proteins could serve as alarmins, in LOs lacking SP-B, the levels of those same cytokines are not present or inducible. Other potentially deleterious cytokines and chemokines (particularly CCL2, CCL20, CXCL8 and CXCL10) and their cellular receptors were dramatically increased. In WT LOs, where SP-B was present, such chemokine levels were held in check (*123*). If infection of the lung epithelia in isolation leads to activation of pro-inflammatory signaling and the production of pro-inflammatory cytokines and chemokines, one could imagine that, when the lung is *in situ* and connected to a vascular system, these molecules would attract both innate and adaptive immune cells from the bloodstream, in turn worsening the situation by creating a positive feedback loop – “cytokine storm” (*124*). The actual mechanism by which surfactant reduces cellular infection is unclear. It may be due to its barrier function, or to the creation of micelles that entrap viral particles, or to its modulatory effects on inflammatory mediators, or to a combination of all these actions (*109, 125–129*). The physical-chemical modes-of-action need to be investigated further but are beyond the scope of this study.

In a patient, persistent viral infection can lead to surfactant depletion if consumption outstrips production, enabling viral spread and inflammatory modes of cell death (*5, 6*). It is under those circumstances that *CARDS* resembles *neonatal RDS* as a surfactant-deficient respiratory failure. The translational question would be whether aggressive and timely surfactant repletion (and/or SP-B replacement) could be therapeutic clinically for restoring lung function following SARS-CoV-2, not just through its physical-chemical attributes but by also inhibiting viral dissemination, modulating local immune responses and inflammatory cascades, preventing chronic inflammation, and reducing cell death -- as we demonstrated above in the LOs using surfactant-based therapeutics. Anecdotal clinical use of surfactants to date has not been systematically timed, adequately controlled, or sufficiently statistically powered (*130–132*).

It is also well-known that COVID-19 hits deprived socio-economic populations and certain racial groups with increased incidence and severity (*133*). Other than a greater prevalence of co-morbidities in those groups (e.g., obesity, diabetes, chronic obstructive lung disease, occupational exposures), neither pre-clinical models nor clinical trials to date have offered much insight into the causes of this disparity (*134*) or how to approach it therapeutically. A handful of therapeutics have been approved to treat COVID-19, reduce its severity, or serve as a pre-exposure prophylaxis (*135–137*). But with emergent SARS-CoV-2 variants, these therapeutics may be less effective (*138*), and clinical trials testing new drugs for new variants in a diverse group of individuals may not be feasible. Our hiPSC derived lung organoid platform is thus unique in that we can differentiate hiPSCs from various backgrounds and sexes into LOs to determine differences in responses to infection and therapeutic efficacy with different compounds. While such a detailed exploration is beyond the scope of this study, the patient-specific hiPSC-derived LOs we used in this work from men and women of diverse racial groups did yield some interesting observations that should be pursued in greater depth using experimental systems like ours **[Supplementary Fig. 5]**.

Our transcriptional interrogation of the infected LOs showed DEGs consistent with those observed following infection of primary human lung cells (*26*). Although this model system lacked immune and vascular cells, the response of the epithelial and fibroblast cells to acute infection revealed immune specific changes. Our scRNAseq data revealed temporal viral transcription and support mechanisms of viral entry that are likely independent of ACE2, TMPRSS2, and FURIN but dependent on other entry routes mediated by cathepsin B/L/S and CD147/BSG. Late endocytosis of virus into cells not expressing ACE2 is clearly a potent entry route (as demonstrated by our Apilimod assay **[Fig. 2l]**) which was nevertheless successfully suppressed by SP-B and surfactants. BSG is a constitutively expressed receptor in epithelial cells and is also a putative receptor for HIV-1 and measles (*139*). Acute infection of human lung induces rapid changes in innate immune responses via interferon signaling, oxidative stress, HIF signaling, and cellular senescence. The variability in the incidence of infection and severe disease from SARS-CoV-2 among neonates, children, and adults is considerable. ACE2 dynamically changes with age, increasing in late adulthood (*120, 140*) and is increased in adults with co-morbidities including smoking (*141–143*). The influence of age, inflammation, and other co-morbidities on the expression of non-canonical viral entry receptors unveiled in this study will require further investigation.

## MATERIALS AND METHODS

### EXPERIMENTAL MODEL AND SUBJECT DETAILS

#### Cell Lines

Vero E6, A549, SK-LU-1, Caco2, and Calu-3 cell lines were obtained from ATCC. Huh7.5 cells were obtained from Apath LLC. TMPRSS2-VeroE6 were obtained from Sekisui XenoTech. A549, SK-LU-1, Caco2, and Calu-3 cells were propagated in MEM (Corning), 10% FBS, Penicillin-Streptomycin (Gibco). Vero E6, TMPRSS2-VeroE6, and Huh7.5 cells were propagated in DMEM (Corning) with 10% FBS and Penicillin-Streptomycin (Gibco) with Geneticin added for TMPRSS2-VeroE6. A549 cells were transduced with hACE2 lentivirus (ASMBio, Lot#200723) or firefly luciferase lentivirus (AMSBio, Lot #200325) with an MOI of 8 with 8mg/mL of polybrene. Transduced cells were selected with 1mg/mL puromycin for 3 days.

Human Embryonic Stem cell and human induced pluripotent stem cell lines were used in accordance with guidelines provided by the National Institutes of Health (NIH). Supplementary table 1 highlights the human iPSCs used and how they were derived. The SP-B mutant lines were previously published (*20*). These cell lines have been validated by immunofluorescence for markers of pluripotency, karyotype and genomic analysis if applicable. The hiPSCs and the human embryonic stem cell line H9 (WiCell), were cultured on matrigel coated (Matrigel GFR; Corning, #354230) plates in mTeSR medium (StemCell Technologies #85850). Medium was changed daily, and cells were passaged using ReLeSR (StemCell Technologies #05872) every 5–7 days. Cultures were maintained in an undifferentiated state in a 5% CO2/ambient air environment.

Authentication of Cell Lines is based on guidelines from the International Cell Line Authentication Committee (iclac.org): Authentication testing was performed on established cell lines regardless of the application. The pluripotency of the ESC and iPSC cell lines were confirmed through immunofluorescence for pluripotent markers (Oct4, Nanog, TRA1-81, SOX2, TRA 1-60) and normal karyotype was also confirmed prior to use. Early passages at the time of verification of pluripotency and karyotype are cryopreserved and those stocks utilized for further experimentation.

Different cell lines and derivatives were never manipulated together to avoid the possibility of cross-contamination. Furthermore, all cell lines were used at early passage. Upon reaching higher passage number, cells were discarded, and cultures restarted from early passage cryovials. Monthly screening for mycoplasma using the MyoSEQ ThermoFisher Mycoplasma Detection kit was performed, and new cells/cell lines were maintained in quarantine until confirmed to be mycoplasma negative.

Primary normal human bronchial epithelial cells (NHBECs) were sourced from Lonza (NHBE CC-2540; Walkersville, MD). These Lonza NHBECs were obtained from a 65-year-old Caucasian male without identifiers. The NHBECs provided were tested and certified negative for HIV, HBV, HCV, sterility, and mycoplasma. The NHBECs were expanded with PneumaCult^TM^ Ex-Plus media (StemCell #05040, Tukwila, WA). Cells were seeded at 5×10^6^ cells/ T75 flask and incubated at 37°C, 5% CO_2_. Culture media was changed every other day. Once cells reached 70-80% confluency, they were dissociated using Animal Component-Free Cell Dissociation kit (StemCell #05426, Tukwila, WA). When cells were ∼70-80% confluent, they were dissociated and seeded on collagen-coated 6.5 mm transwells (Corning #29442-082, VWR). Transwells were pre-coated with 50µg/mL Collagen type I from Rat tail (BD Biosciences #354236) at 7.5 µg/cm^2^. Collagen-coated transwells were rehydrated with 100 µL PneumaCult^TM^ Ex-Plus media at 37°C, 5% CO_2_ for 30 min prior to seeding NHBECs for ALI culture. NHBECs were then seeded in the apical chamber at 5×10^4^ cells in 100uL of media, for a total of 5×10^4^ cells/ 100 µL of media per collagen-coated transwell with 500µL of media in the basolateral chamber. The cells were then incubated at 37°C, 5% CO_2_ overnight. On Day 1, the media in the apical and basolateral chambers were exchanged with 100 µL and 500 µL, respectively, with fresh PneumaCult^TM^ Ex-Plus media. NHBECs were 80-100% confluent the day after seeding. The next PneumaCult^TM^ Ex-Plus media change was on Day 3. On Day 4-7, or until culture reached confluency, the apical and basolateral chambers were fed PneumaCult^TM^ ALI media (StemCell #05021) daily supplemented with 10µM ROCK inhibitor (Tocris Y-27632). On Day 8, the apical media was removed, and the basal media replaced with PneumaCult^TM^ ALI media without Y-27632. Subsequent media changes were every other day thereafter. On Day 14 post-airlift, the apical surfaces were washed with DPBS, once per week. Cells were grown in 37°C, 5% CO_2_ incubator until four weeks post airlift.

#### Directed differentiation of hESC and hiPSCs to lung organoids

For human iPSC-lung organoid generation, we used H9 embryonic stem cells (WiCell), and 6 iPSC cell lines **[Supplementary Table 1]** which were cultured in feeder free conditions upon Matrigel (Corning #354230) coated plates in mTeSR Plus medium (StemCellTech #100-0276). Media was changed daily, and stem cells were passaged weekly using enzyme free dissociation reagent ReLeSR™ (Stem Cell Tech #05872). Cultures were maintained in an undifferentiated state, in a 5% CO_2_ incubator at 37°C. For lung progenitor spheroid generation, human PSCs were dissociated into single cells with exposure to 20 min of accutase, and approximately 2.0 x 10^5^ cells were seeded on Matrigel-coated plates in mTeSR supplemented with 10μM Y-27632 and incubated for 24 hours. Each iPSC cell line required seeding density optimization with the goal of 50-70% confluency the day after seeding. Definitive Endoderm (DE) induction medium (RPMI1640 + Glutamax, 2% B27 without retinoic acid supplement, 1% HEPES, 50 U/mL penicillin/streptomycin) supplemented with 100 ng/mL human activin A (R&D), 5 µM CHIR99021 (Stemgent), was added on day 1. On days 2 and 3 cells were cultured in DE media with only 100 ng/mL human activin A supplemented. Anterior Foregut Endoderm (AFE) was generated with serum free basal media (SFBM) made up of 3 parts IMDM:1-part F12, 1% B27 without retinoic acid supplement, 0.5% N2 supplements, 50 U/mL penicillin/streptomycin, 0.25% BSA, 50 µg/mL L-ascorbic acid, 0.45 mM monothioglycerol. SFBM was supplemented with 10 µM SB431542 (R&D) and 2 µM Dorsomorphin (StemGent) on days 4-6. On day 7, AFE cells were dissociated with accutase exposure for 10 min and 3 x 10^5^ cells were passaged as aggregates into 150 µL of cold matrigel droplets to make Lung Progenitor Cell (LPC) Spheroids. The matrigel containing cells were polymerized for 30-60 min in a 37°C incubator and then LPC media (SFBM supplemented with 10 ng/mL of human recombinant BMP4, 0.1 µM of retinoic acid (RA), 3 µM of GSK3β inhibitor/Wnt activator CHIR99021 and 10 µM of Rock Inhibitor Y-27632) was added to submerge the matrigel droplets. LPC media was changed every other day for 9-11 days without Y-27632. Analysis of the surface antigen CPM, or the intracellular marker NKX2-1 should be performed at the end of this differentiation period to determine the efficacy of differentiation. If the LPC spheroids express > 50% CPM/NKX2-1, lung organoid differentiation may proceed.

To generate lung organoids (proximal, distal, and whole lung), LPC spheroids were incubated in 2 U/ml dispase for 30 min at 37°C with manual pipetting every 15 min to break apart the matrigel. Cold PBS was added, transferred to a 15 mL conical tube, and then centrifuged at 400 x g for 5 min. The supernatant was carefully removed then 3 mL of cold PBS was added to the pellet, mixed, and then centrifuged again at 400 x g for 5 min. The supernatant was carefully removed, and the pellet resuspended in 2-3 mL of TrypLE Express (Gibco # 12605010) for 12 min at 37°C to keep the LPC spheroids as aggregates. The reaction was quenched with 2% FBS in DMEM/F12 then centrifuged at 400 x g for 5 min. The supernatant was removed, and the cell pellet resuspended in 1ml of quenching media supplemented with 10 µM Rock inhibitor (Y-27632) and a cell count was performed. Cells were embedded into cold matrigel as aggregates at 5 × 10^4^ cells per 200 µL of Matrigel and transferred into the apical portion of 0.4 µm translucent 12-well inserts (VWR 10769-208).

To generate 3D human proximal lung organoids, we modified a previously published protocol. After the LPC spheroids were passaged onto transwells, Proximal lung organoid media was added to the basal side of the insert. The medium was made up of SFBM supplemented with 250 ng/mL FGF2, 100 ng/mL rhFGF10, 50 nM dexamethasone (Dex), 100 µM 8-Bromoadenosine 3’,5’-cyclic monophosphate sodium salt (Br-cAMP), 100 µM 3-Isobutyl-1-methylxanthine (IBMX) and 10 µM ROCK inhibitor (Y-27632). Proximal lung organoid media was changed every other day for 2-3 weeks.

To generate 3D human distal lung organoids, we modified a previously published protocol. After the LPC spheroids were passaged onto transwells, Distal lung organoid media was added to the basal side of the insert. Distal lung organoid maturation media was made up of distal basal media (95% F12 media, 0.25% BSA, 15 mM HEPES, 0.8 mM calcium chloride (Wako), 0.1% ITS premix (Corning), 1% B27 without RA and 50 U/mL penicillin-streptomycin) supplemented with 3 µM CHIR99021, 10 µM SB431542, 10 ng/ml FGF7, 0.1 µM RA, 50nM Dex, 100 µM Br-cAMP, 100 µM IBMX and 10 µM ROCK inhibitor (Y-27632). Distal lung organoid media was changed every other day for 2-3 weeks without Y-27632.

To generate 3D human whole lung organoids, we modified a previously published protocol. LPCs in matrigel droplets were passaged as above. Whole lung organoid induction media was added to the basal side of the transwell with SFBM supplemented with 10 ng/mL FGF7, 10 ng/mL FGF10, 10 ng/mL EGF, 3 µM CHIR99021, and 10 µM ROCK inhibitor (Y-27632). Media changes without Y-27632 were done every other day for 6 days. Whole lung organoid branching medium was added using SFBM supplemented with 10 ng/mL FGF7, 10 ng/mL FGF10, 10 ng/mL EGF, 3 µM CHIR99021, 0.1 µM retinoic acid and 10 ng/mL VEGF/PIGF for 6 days, with media changes every other day. Finally, whole lung organoid maturation medium was added using SFBM supplemented with 10 ng/mL FGF7, 10 ng/mL FGF10, 10 ng/mL EGF, 3 µM CHIR99021, 0.1 µM All-trans retinoic acid, 10 ng/mL VEGF/PIGF, 50 nM Dex, 100 µM Br-cAMP, and 100 µM IBMX for 6 days, with media changes every other day.

For infections in intact 3D lung organoids, the matrigel droplets were washed with PBS, then 1 ml of Cell Recovery Solution (Corning #354253) was added per 200 µL Matrigel drops. Matrigel was mechanically dissociated using wide bore P1000 tips, titurating gently to maintain the integrity of the organoids. The mixture was placed in 4°C for 1 h, with manual resuspension every 15 min. After 1 hour, the sample was transferred to a 15 mL conical tube, using a wide bore pipette. The organoids were centrifuged at 200 x g for 1 min. Chilled PBS was added to the pellet and gently mixed then centrifuged at 200 x g for 1 min. The pellet was resuspended in 500 µL of lung organoid media supplemented with 10 µM Y-27632 in 24 well ultra low attachment (ULA) plate (Corning #3473) at approximately 20-30 organoids per well. On the day of infection, the 3D organoids were placed in lung organoid medium without DCI.

For infections in 2D monolayers, 3D lung organoids were dissociated into single cells and seeded at 2×10^4^ cells per well of a matrigel coated 96-well plate approximately 3 days before infection. Matrigel containing lung organoids was incubated in 2 U/ml dispase for 30 min at 37 °C with manual pipetting every 15 min to break apart the matrigel. Cold PBS was added to the well, transferred to a 15 mL conical tube and then centrifuged at 400 x g for 5 min. The supernatant was carefully removed then 3 mL of cold PBS was added to the pellet, resuspended, and then centrifuged again at 400 x g for 5 min. The supernatant was carefully removed, and the pellet resuspended in 2-3 mL of TrypLE Express (Gibco # 12605010) for 15-20 min at 37°C to dissociate the organoids into single cells. The reaction was quenched with 2% FBS in DMEM/F12 then centrifuged at 400 x g for 5 min. The supernatant was removed, and the cell pellet resuspended in 1ml of quenching media supplemented with 10 µM Rock inhibitor (Y-27632) and a cell count was performed. Cold PBS was added to the mixture then centrifuged at 400 x g for 5 min. Supernatant was carefully removed and resuspended in 2-3 mL of TrypLE Express (Gibco # 12605010) for 20 min at 37 °C. Reaction was quenched with 2% FBS in DMEM/F12 then centrifuged at 400 x g for 5 min. The supernatant was aspirated, and the cell pellet resuspended in 1 mL of quenching media supplemented with 10 µM Rock inhibitor (Y-27632). Cell count was performed, and 2 x 10^4^ cells were seeded per well of a matrigel coated 96-well plate with 100 µL of respective lung organoids media. The wells were 90-100% confluent prior to infections. On the day of infection, the media was replaced with lung organoid medium without DCI.

All materials used can be found in **Supplementary Table 3**. All media recipes can be found in **Supplementary Table 4.**

#### Infection by Pseudotyped virus

Vesicular Stomatitis Virus (VSV) pseudotyped with spike proteins of SARS-CoV-2 were generated according to a published protocol (*46*). Briefly, HEK293T, transfected to express the SARS-CoV-2 spike protein, were inoculated with VSV-G pseudotyped ΔG-luciferase or GFP VSV (Kerafast, MA). After 2 h incubation at 37°C, the inoculum was removed and DMEM supplemented with 10% FBS, 50 U/mL penicillin, 50 mg/mL streptomycin, and VSV-G antibody (I1, mouse hybridoma supernatant from CRL-2700; ATCC). Pseudotyped particles were collected 24 h post-inoculation, centrifuged at 1,320 x g and stored at -80°C until use.

For pseudovirus-GFP infections, VSV-G pseudotyped ΔG-GFP was added to 3D or 2D hiPSC derived lung organoids at an MOI of 1-5 for 24 h. After infection, the cells were washed with 1× Dulbecco’s phosphate-buffered saline (PBS) three times. The 3D organoids were prepped for flow cytometry/FACS or single cell RNA sequencing per the protocol below. The monolayer infected lung cells were washed with PBS three times then fixed in 4% PFA for immunohistochemistry (see below).

#### SARS-CoV-2 viruses

All work with SARS-CoV-2 was conducted in Biosafety Level-3 conditions at the University of California San Diego following the guidelines approved by the Institutional Biosafety Committee.

##### Isolates

SARS-CoV-2 isolates WA1 (USA-WA1/2020, A), B.1 (USA/NY-PV08410/2020, B.1, D614G) and Beta (hCoV-19/South Africa/KRISP-K005325/2020, B.1.351, Beta) were acquired from BEI and passaged once through primary human bronchial epithelial cells (NHBECs) differentiated at air-liquid interface (ALI) to select against furin site mutations. Virus was then expanded by one passage through TMPRSS2-Vero cells.

Alpha (hCoV-19/USA/CA_UCSD_5574/2020, B.1.1.7, Alpha) was isolated from a patient nasopharyngeal swab under UC San Diego IRB #160524. The original clinical isolate sequence can be accessed at GISAID accession name: EPI_ISL_751801. Material from nasopharyngeal swab in PBS with 1x Antibiotic-Antimycotic (Thermo Fisher Scientific #15240062) was added to the apical chamber of 4 weeks old ALI NHBECs after two 30 min PBS washes at 37°C. 1 x Antibiotic-Antimycotic was also added to the media in basal chambers. After 24 h, input was removed, and apical surfaces were washed. Apical washes (passage 1) were taken daily and stored at -80. Fresh NHBECs at ALI were inoculated with passage 1 virus as above to produce passage 2 and with passage 2 to produce passage 3. Titers were expanded by infecting TMPRSS2-Vero with passage 3 virus. Gamma (hCoV-19/Japan/TY7-503/2021, P.1, Gamma), and Delta (hCoV-19/USA/PHC658/2021, B.1.617.2, Delta) were acquired from BEI and expanded on TMPRSS2-VeroE6 cells.

Omicron (hCoV-19/USA/CA-SEARCH-59467/2021, BA.1, Omicron) was isolated from a patient sample under UC San Diego IRB #160524. Original patient isolate sequence can be found at GISAID (EPI_ISL_8186377). Serial dilutions in DMEM + 3% FBS were made from a positive nasopharyngeal swab stored in viral transport media. Dilutions were incubated on TMPRSS2-VeroE6 cells and monitored for CPE. When CPE was observed, the contents were transferred to fresh cells for a total of 3 passages on TMPRSS2-VeroE6 cells.

For single cell RNAseq of the mixed organoids, SARS-CoV-2 isolate WA1 (USA-WA1/2020, A) was acquired from BEI and propagated on VeroE6 cells resulting in mutation of the furin cleavage site (*144, 145*).

All viral stocks were clarified and stored at -80°C. Titers were determined by plaque assay or fluorescent focus assay on TMPRSS2-VeroE6 cells. All stocks are verified by full genome sequencing. Original sample sequencing links can be found in **Supplementary Table 5.**

#### Infection of NHBECs at ALI

After two 30 min incubations with PBS at 37°C, 5% CO_2_, virus diluted in PBS was added to the apical chamber in 100 µL. Virus was removed after 24 h and apical washes (150 µL PBS with 10 min incubation at 37°C, 5% CO_2_,) were taken daily and stored at -80°C. Titer was determined by fluorescent focus assay on TMPRSS2-Vero cells. Infected transwells were fixed in 4% PFA for immunofluorescence staining.

#### Plaque assay

Ten-fold serial dilutions were made in DMEM and added to 12-well plates of confluent VeroE6 cells for 1 h at 37°C with occasional rocking. Input was removed and monolayers were overlaid with 1mL of a 1:1 mixture of 1.2% agarose and 2 x MEM complete (2 x MEM supplemented with 8% FBS, 2 x L-glutamine, 2 x non-essential amino acids, and 2 x sodium bicarbonate). Plates were incubated 48 h at 37°C and then fixed with 2 mL 10% formaldehyde in PBS for 24 h at RT. Overlays were removed, and plaques were visualized by staining with 0.025% crystal violet in 2% EtOH. Plaque assays were performed and counted by a blinded experimenter. Plaque assays were counted and recorded by a blinded observer.

#### Fluorescent focus assay

Ten-fold serial dilutions were made in DMEM and added to 96-well plates of confluent TMPRSS2-VeroE6 cells. After 1 h at 37°C, input was removed, and wells were overlaid with 100 µL 1% methylcellulose in DMEM + 1% FBS and 1 x Pen/Strep. Plates were fixed after 24 h incubation at 37°C by addition of 150 µL 8% formaldehyde for at least 30 min at RT. Cells were stained with anti-nucleocapsid primary antibody (GeneTex, gtx135357) and anti-rabbit AlexaFluor secondary (Thermo Fisher Scientific) and plates were imaged on the Incucyte S3 imager. Foci were counted using the Incucyte onboard software tools.

#### Virus quantification, FFU immunofluorescence imaging and analysis

3D hPSC-LOs were removed from matrigel intact as described above. They were plated in 24 well ultra-low attachment plates in lung organoid base media, without DCI, at a concentration of 20 organoids per well. hPSC-LOs that were dissociated and plated as a monolayer were also changed to lung organoid base media without DCI, prior to infection. 3D LOs were infected with SARS-CoV-2 at an MOI of 1-3 and incubated for 24 -72 h at 37 °C. Dissociated LOs were infected with SARS-CoV-2 at an MOI of 0.1.

For the compound experiments, 3D lung organoids in suspension were pretreated with DMSO (control), cathepsin inhibitors ONO5334 and Apilimod and remdesivir (all at a concentration of 5 µM). Two hours after the addition of compounds, SARS-CoV-2 was added for 24 h. Mock infections were performed using viral growth media (DMEM + 3% FBS). Infected and mock controls were washed three times in PBS and lysed in radioimmunoprecipitation assay (RIPA) lysis buffer (Millipore 20-188) at 2 x with protease inhibitor for western blot analysis.

For infections with poractant alfa and recombinant SP-B, hiPSC lung organoids were dissociated & plated as a monolayer (4 x 10^4^ cells/well) (along with VeroE6 cells as a positive control), pre-incubated with exogenous surfactant (poractant alfa [Curosurf] 1 mg/ml) or with recombinant SP-B [rSP-B] (5 µM) for 30 min prior to infection with SARS-CoV-2 at an MOI of 0.1. After 1 hour of infection, cells were overlayed in media supplemented with 1.2% carboxymethylcellulose as discussed below.

For infections of VeroE6 cells using carboxymethylcellulose, media was prepared using 500 mL lung organoid base media or DMEM-glutamax supplemented with 3% FCS, 2% gentamycin, 1% penicillin/streptomycin plus 6 g autoclaved carboxy methyl cellulose. iPS-derived human lung organoids were dissociated and plated as a monolayer then infected with SARS-CoV-2 at the indicated MOI. After 1 h of infection, cells were overlayed in carboxymethylcellulose media. After incubation for up to 72 h, overlay was removed and cells were fixed in 4% formaldehyde for 30 min at RT. Fixed cells were washed in BD CytoPerm, and then stained overnight with a primary nucleocapsid antibody (GeneTex GTX135357), counterstained with an anti-rabbit AF647 secondary antibody (ThermoFisher), and nuclei counterstained with 1 µM Hoechst or 1 µM Sytox Green.

Whole well scans were captured with an Incucyte S3 or 10 x 0.45 NA objective using a Nikon Ti2-E microscope with a Qi-2 camera and Nikon Elements 5.02.02 acquisition software utilizing the Jobs module. Wells were imaged for red, green and/or far-red fluorescence using a SpectraX light engine (Lumencor) with individual LFOV filter cubes (Semrock) (excitation/emission maxima at 554/609, 470/525 and 618/698 nm, respectively). FFU analysis was carried out using a custom macro written in the Fiji distribution of ImageJ. The macro leverages machine learning using Ilastik (*146*), StarDist (arXiv:1806.03535v2) and GPU acceleration using CLIJ to process data files (*147*). Nuclei were segmented using StarDist, and positive cell stain masks generated using a trained model in Ilastik. Masks were used to count the number of nuclei in clusters, and to define the staining intensity of the positive clusters, and morphological characteristics and size of FFU, and the distribution of the clusters within the well and from each other. The cluster shape was also assessed by generating a ratio of the perimeter and convex hull of each cluster, with values approaching 1 being circular and values approaching zero becoming more irregular.

#### FFUTrackR data visualization

Image J datalogs from analyses of whole well scans were visualized using the R Shiny app FFUTrackR (https://croker.shinyapps.io/ffutrackr). The FFUTrackR pipeline was developed using Shiny in R 3.6.3., the R packages Tidyverse and plotrix for basic data loading and filtering utilities. shinydashboardPlus and shinyWidgets are used for module management and custom input controls. FFUTrackR requires R packages dplyr, ggpubr, and ggbeeswarm for basic data handling and plotting utilities. The code for FFUTrackR is available at: https://github.com/WeiqiPeng0/FFU.TrackR.

#### Virus-induced cell death

SARS-CoV-2-triggered cell death of Vero E6 cells was monitored using an Incucyte S3. Virus was added to Vero E6 cells at the indicated MOI for 1 h in DMEM/10% FBS at 37°C/5%CO_2_, then BH3 mimetics added for up to 48 h. Cell viability was monitored using 1 µg/mL propidium iodide. Five fields of view at 10x magnification representing 33.6% well coverage was monitored for changes in cell viability every 6 h. PI-positive cells were identified using Incucyte software.

#### Bulk RNA sequencing preparation and analysis

RNA was purified from TRIzol lysates using Direct-zol RNA Microprep kits (Zymo Research R2061) according to manufacturer recommendations that included DNase treatment. RNAseq assay was performed using an Illumina NEXTSeq 500 platform in the SBP Genomics Core. Briefly, PolyA RNA was isolated using the NEBNext® Poly(A) mRNA Magnetic Isolation Module and barcoded libraries were made using the NEBNext® Ultra II™ Directional RNA Library Prep Kit for Illumina® (NEB, Ipswich MA). Libraries were pooled and single end sequenced (1X75) on the Illumina NextSeq 500 using the High output V2 kit (Illumina Inc., San Diego CA).

For viral RNA analysis, sequencing reads were aligned to the SARS-CoV-2/human/USA/WA-CDC-WA1/2020 genome (GenBank: MN985325.1) using Bowtie2 and were visualized using IGV software. After further filtering and quality control, the R package edgeR29 was used to calculate reads per kilobase of transcript per million mapped reads (RPKM) and log2 [counts per million] matrices as well as to perform differential expression analysis. PCA was performed using log2 [counts per million] values and gene set analysis was run with WebGestalt30. Heatmaps and bar plots were generated using Graphpad Prism software, v.7.0d. In the volcano plots, differentially expressed genes (P-adjusted value < 0.05) with a log2 [fold change] > 1 are indicated. Read data was processed in BaseSpace (basespace.illumina.com). Reads were aligned to Homo sapiens genome (hg19) using STAR aligner (https://code.google.com/p/rna-star/) with default settings.

We used Cutadapt v2.3 [147] to trim Illumina Truseq adapters, polyA, and polyT sequences. Trimmed reads were first aligned to SARS-CoV-2 genome version NC_045512v2 using STAR aligner v2.7.0d_0221 [148] with parameters according to ENCODE long RNA-seq pipeline (https://github.com/ENCODE-DCC/long-rna-seq-pipeline). Unaligned reads to SARS-CoV-2 genome were subsequently mapped to human genome hg38 using STAR and parameters as above. Gene expression levels were quantified using RSEM v1.3.1 [149] and Gencode gene annotations v32 (Ensembl 98). RNA-seq sequence, alignment, and quantification qualities were assessed using FastQC v0.11.5 (https://www.bioinformatics.babraham.ac.uk/projects/fastqc/) and MultiQC v1.8 [150]. Biological replicate concordance was assessed using principal component analysis (PCA) and pairwise Pearson correlation analysis. Lowly expressed genes were filtered out by applying the following criterion: estimated counts (from RSEM) ≥ number of samples * 5. Filtered estimated read counts from RSEM were compared using the R Bioconductor package DESeq2 v1.22.2 based on generalized linear model and negative binomial distribution [151]. Genes with Benjamini-Hochberg corrected p-value < 0.05 and fold change ≥ 2.0 or ≤ -2.0 were selected as differentially expressed genes. Differentially expressed genes were then analyzed using Ingenuity Pathway Analysis (Qiagen, Redwood City, USA) and Gene Set Enrichment Analysis (GSEA) (*148*).

#### Single-cell lung organoid preparation for scRNA-seq

3D hPSC-LOs in ultra-low attachment plates were dissociated into single cells using accutase (Gibco) at 37 °C for 10 min, 24hpi. The dissociated organoids were pelleted and resuspended with DMEM F12 + 10% FBS. The resuspended organoids were then placed through a 40 µm filter to obtain a single-cell suspension. To remove the dead cells, 1 µL Dead Cell Removal cocktail (Annexin V, STEMCELL Technology) was added into the dissociated CO/PCCO single cells suspension together with 1 µL Biotin selection cocktail (Annexin V, STEMCELL Technology), gently mixed and incubated at RT for 3 min. After 3 min incubation, RapidSpheresTM (STEMCELL Technology) was vortexed for 30 s at RT and 2 µL added to the above mixture. At the same time, 850 µL lung organoid base medium was added and gently pipetted 2∼3 times. Tubes were then placed into a magnetic holder and incubated for 3 min at RT. After incubation, the supernatant was collected in a new tube for 10X GEM generation. For each of the SC-RNA-seq library, we pooled 20 individual LOs derived from H9 cells and iPSC-lines specified in the manuscript.

For the hPSC lung organoids outside the BSL3, they were dissociated from matrigel as described above using dispase and trypsin. The resuspended organoids were then placed through a 40 µm filter to obtain a single-cell suspension and stained with DAPI followed by sorting of live cells using the FACS Aria Fusion Flow Cytometer (BD Biosciences). The sorted cells were washed with 1× PBS + 0.04% BSA, counted manually after sorting and the cell viability was assessed with trypan blue.

#### Single cell library preparation and sequencing

10X sc-RNA-seq-3’-V3.1 kit (10X Genomics) was used to generate the GEM, cDNA and sequencing library according to the manufacturer’s instructions (10X Genomics). Briefly, live cells were partitioned into nanoliter-scale Gel Bead-In-Emulsions (GEMs) with the 10x Chromium Controller (10X Genomics), 1000 cells were targeted. Upon cell lysis and dissolution of the Single Cell 3′-V3.1 Gel Bead within the droplet, primers containing an Illumina P7 and R2 sequence, a 14 bp 10XBarcode, a 10 bp randomer, and a poly-dT primer sequence were released and mixed with the cell lysates and bead-derived Master Mix. Barcoded, full-length cDNA from poly-adenylated mRNA was then generated in each individual bead, then individual droplets were broken and homogenized before the remaining non-cDNA components were removed with silane magnetic beads (Invitrogen). The libraries were then size-selected, and the R2, P5 and P7 sequences were added to each selected cDNA during end repair and adapter ligation. After Illumina bridge amplification of cDNA, each library was sequenced with the PE100 reads on the Novaseq6000 at the IGM Core in UCSD (mixed lung organoid samples), and The Scripps Oceanography Genomics Core (proximal lung organoid samples), to a depth of approximately 120M reads per sample.

The FASTQ files were imported to a 10x Cell Ranger data analysis pipeline (v.3.0.2) to align reads, generate feature–barcode matrices and perform clustering and gene expression analysis. In a first step, cellranger mkfastq demultiplexed samples and generated. fastq files; and in the second step, cellranger count aligned fastq files to the reference genome and extracted gene expression unique molecular identifier (UMI) count matrix. To measure viral gene expression in lung organoids, we built a custom reference genome by integrating the four virus genes and GFP into the 10x pre-built human reference (GRCh38 v.3.0.0) using cellranger mkref. The sequences of four pseudo-viral genes (VSV-N, VSV-P, VSV-M and VSV-L) and SARS-CoV-2 genes (E, M, N, S, ORF1ab, ORF3a, ORF6, ORF7a, ORF8, and ORF10) were retrieved from NCBI https://www.ncbi.nlm.nih.gov/nuccore/335873).

#### scRNA-seq data analysis for lung organoids

We prepared a combined human genome version hg38 and SARS-CoV-2 genome version NC_045512v2 for alignment of 10X scRNA-seq raw data,. We used human Gencode gene annotations v32 (Ensembl 98) augmented with SARS-CoV-2 predicted genes from NCBI for the Cell Ranger processing step. scRNA-seq reads from each sample were aligned to the combined genome using Cell Ranger v5.0.1. Data QC, filtering, integration, clustering, and differential expression were performed using Seurat v4.0.5 (*149*) and Harmony (*150*). Prior to integration of samples, cells with <40% mitochondrial content and < 98 percentile of genes detected (to remove potential doublets/multiplets) were kept in each sample for downstream analysis. The samples were merged and gene counts for the merged dataset were normalized using *NormalizeData()*. 3000 variable features were determined using *FindvariableFeatures()* and the data scaled using *ScaleData()*. PCA components were computed using *RunPCA()* and subsequently the samples were integrated using *RunHarmony()* function. Clusters of cells were computed using *RunUMAP(reduction=”harmony”, dims=1:40)*, *FindNeighbors(reduction=”harmony”, dims=1:40)*, and *FindClusters(resolution=0.3).* Cluster markers were found using *FindAllMarkers().* Infection and IFNome scores were computed using *AddModuleScore().* Differential expression analyses were performed using *FindMarkers()*. Feature, violin, and other plots were prepared using Seurat or ggplot2. Pathway analyses of differentially expressed genes were performed using Ingenuity Pathway Analysis (Qiagen, Redwood City, USA) and Gene Set Enrichment Analysis (GSEA) (*148*).

To obtain a list of top interferon-stimulated genes (ISGs), all human type I interferon datasets were taken from the Interferome v2.0 database (*96*). For each dataset, significantly up-regulated genes (log2 fold change > 1; unadjusted p-value < 0.05) were ranked by the magnitude of their log2 fold changes. These ranks were then combined across datasets by geometric mean using the TopKLists (*151*) R package to produce an overall ranked list of type I ISGs (Interferon Stimulated Genes) identified.

#### Identifying & Labeling Endosomes (Macropinosomes)

Endosomes (specifically macropinosomes) were labeled and quantitated utilizing previously established procedures (*152–154*). Macropinosomes in live cells were labelled by their uptake of high molecular weight fluorescent dextran (FITC-Dextran). In brief, live cells are incubated with 1 mg/mL high molecular weight FITC-Dextran for 30 min, washed with cold PBS, fixed with 3.7% paraformaldehyde and nuclei stained with DAPI. Quantitation was as previously described using image analysis facilitated by Cytation 5 automated imaging and the Gen 5 software.

Macropinocytosis in lung cells is also dependent on PI3K; uptake in these cells was inhibited by treatment with LY294002 (25 mM), an inhibitor of PI3K. Cells were pre-treated with the respective inhibitors for 0.5-3 hours prior to infection. At least 25,000 cells-per condition were analyzed for n=3 replicates.

#### Western blot

Cell samples were washed 3 times with PBS to remove traces of media components, pelleted and lysed with ice cold RIPA buffer containing protease and phosphatase inhibitors (Thermoscientific Halt kit). Samples were vigorously vortexed and centrifuged at 16,000 x g for 25 min at 4°C to separate the proteinous supernatant from the cellular debris. Protein concentration of the supernatant was calculated using a BCA assay kit (ThermoScientific, 23225). 10 µg of protein and Nupage sample buffer with diothiothreitol was boiled at 95°C for 3 min then loaded into a bis-tris gel and run with MES running buffer. The gel was transferred to 0.45 µm PVDF membrane for 60 min at 20V, then blocked in 5% w/v nonfat dry milk (NFDM) in tris buffered saline (TBS) for 60 min. Primary antibodies were diluted in 1% NFDM in TBS with 0.1% Tween-20 (TBST) and incubated overnight at 4°C. The membranes were then washed in TBST 3 times for 5 min on an orbital shaker. Secondary antibodies (LICOR goat anti rabbit 800CW and goat anti-mouse 680, 1:10,000) were diluted in 1% NFDM for 60 min then rinsed with TBST before imaging on the odyssey near infrared scanner.

#### Immunofluorescence

Dissociated lung cells were washed twice in PBS, then fixed in BD Fix/Perm containing 4% formaldehyde for 30 min at RT. Fixed cells were washed twice in BD Cytofix/Cytoperm and stained for SARS-CoV-2 with a primary Nucleocapsid antibody at 1:2000 (GeneTex GTX135357) conjugated to AF594 (ThermoFisher A20185) and nuclei counterstained with Hoechst or Sytox Green at 1:5000. 3D Organoids were fixed in 4% paraformaldehyde for 1 h, then washed twice with PBS. Fixed organoids were embedded in a 30% w/v solution of sucrose to preserve the fine cytoarchitecture from cryo-damage. Organoids were then imbedded into OTC and sliced at 10 µm. Serial sections were mounted onto glass slides and allowed to air dry before staining. Sections were incubated in primary antibody diluted in 5% v/v donkey serum and 5% w/v bovine serum albumin with TX-100 for 24 h at 4°C then secondary antibodies at room temperature for 1 h. Nuclei were counterstained by DAPI (1:5000). The information for the primary and secondary antibodies is provided in **Supplementary Table 6**. The figures were processed using ImageJ software.

#### Intracellular flow cytometry analysis

Flow cytometry intracellular staining was performed following the instruction manual for the Fixation/Permeabilization Solution Kit (BD Biosciences). In brief, infected cells were washed twice with PBS, then incubated in Zombie UV (BioLegend 423107) per the manufacturer’s recommendations for 30 min at room temperature. Cells were washed in FACS buffer (PBS + 2% FBS) once and resuspended in Fixation/Permeabilization solution at 4 °C for 30 min. The cells were washed twice in 1× Perm/Wash buffer, incubated with primary Nucleocapsid antibody (GeneTex GTX135357) conjugated to AF594 (ThermoFisher A20185) at 4 °C for 30 min in the dark, and washed twice before flow cytometry analysis. Flow cytometry was performed using mock infected LOs as a negative control and gates were set to exclude the dead cells (Zombie UV stained) using the LSR Fortessa X-20 (BD Biosciences). All experiments were done a minimum of three separate times in three technical replicates. Data analysis was performed using FlowJo software and statistical analyses were done in Prism 8 (GraphPad). The information for the primary antibodies and antibodies is provided in **Supplementary Table 6.**

#### Poractant Alfa and recombinant SP-B

Vero cells were plated in a monolayer in a 96 well plate and 3 titrations (1 mg/mL, 0.1 mg/mL and 0.01 mg/mL) of Poractant alfa (Curosurf) were added 60 min prior to infection. The SARS-CoV-2 variants (Alpha, Beta, Gamma, Delta, WA, and NY) were added at 150 FFU and the plate was placed on a shaker for 1 h. The plate was then overlayed with 150 µL of DMEM/Carboxymethylcellulose/FBS mix and placed in the incubator for 24 h. The hPSC derived lung organoids were plated as a monolayer in a 96 well plate as previously described. They were exposed to 1 mg/mL of Poractant Alfa or 6.6 µM of recombinant SP-B (Prospec #PRO-2585) for 60 min prior to infection. SARS-CoV-2 was added at an MOI of 0.1 and the plate was placed on a shaker for 1 h. The plate was then overlayed with 150 µL of SFBM/Carboxymethylcellulose/FBS mix and placed in the incubator for 24 h. After 24h, the plates were washed twice with PBS and fixed and stained as per the immunofluorescence protocol above.

#### Quantification And Statistical Analysis

Statistical analysis was performed with Prism 8 (GraphPad Software). We compared all time course data related to cell death by two-way analysis of variance (ANOVA) followed by a Šidák multiple comparisons test. For the poractant alfa and rSP-B experiments, a one-way ANOVA or one-tailed t-test was used to determine significance. This was followed by a Šidák multiple comparisons test correction.

## List of Supplementary Materials

Figs. S1 to S8

Tables S1 to S6

Movies S1 to S2

## Acknowledgments

We thank the UC San Diego Center for Advanced Laboratory Medicine Clinical Microbiology and Virology Lab and UC San Diego EXCITE for providing clinical samples for viral isolation and SARS-CoV-2 genome sequencing.

The following reagent was deposited by the Centers for Disease Control and Prevention and obtained through BEI Resources, NIAID, NIH: SARS-Related Coronavirus 2, Isolate USA-WA1/2020, NR-52281. The following reagent was obtained through BEI Resources, NIAID, NIH: SARS-Related Coronavirus 2, Isolate New York-PV08410/2020, NR-53514. The following reagent was obtained through BEI Resources, NIAID, NIH: SARS-Related Coronavirus 2, Isolate hCoV-19/South Africa/KRISP-K005325/2020, NR-54009, contributed by Alex Sigal and Tulio de Oliveira. The following reagent was obtained through BEI Resources, NIAID, NIH: SARS-Related Coronavirus 2, Isolate hCoV-19/Japan/TY7-503/2021 (Brazil P.1), NR-54982, contributed by National Institute of Infectious Diseases. The following reagent was obtained through BEI Resources, NIAID, NIH: SARS-Related Coronavirus 2, Isolate hCoV-19/USA/PHC658/2021 (Lineage B.1.617.2; Delta Variant), NR-55611, contributed by Dr. Richard Webby and Dr. Anami Patel. The CRISPR corrected hiPSC lines from the SP-B mutant patient generously provided by Dr. Darrell Kotton.

## Funding

California Institute of Regenerative Medicine (CIRM) DISC2COVID19-12022 (SLL, EYS)

US-BSF grant 2017176 (BAC)

American Asthma Foundation (BAC)

National Institutes of Health grant AI036214 (CC)

NIH/NCI COVID-19 administrative supplement 3R01CA207189-05S1 (CC)

National Institutes of Health SIG grant S10 OD026929 (UC San Diego IGM Genomics Center)

## Author contributions

Conceptualization: SLL, BAC

Methodology: SLL, RNM, EMK, AEC, AA, CC, BAC

Software: BAC

Validation: SLL, CC, BAC

Formal analysis: SLL, RNM, RM, CMG, KMOG, CC, BJ, BAC, EYS

Investigation: SLL, RNM, EMK, AEC, AA, BAG, RN, REY, YPZ, BJ, CMG, KMOG, CC, AFC, BAC

Resources: SLL, AEC, XS, AFC, EYS, BAC

Funding acquisition: SLL, EYS, BAC

Supervision: AEC, JCL, WP, EG, CJN, KL, SA, AM, LJG, PJH, DMS, XS, AFC, EYS, BAC

Writing – original draft: SLL

Writing – review & editing: SLL, BAC, EYS

## Data and materials availability

### Materials Availability

This study did not generate new unique reagents.

### Data and Code Availability

scRNA-seq data is available from the GEO repository database: GSE214762, GSE214752, GSE214770. RNA-seq data is available from the GEO repository database: GSE214482.

The custom-scripted macro used for automated image analysis of FFU imaging data is available at https://figshare.com/s/13587b74c09c251b3345. (DOI for when we publish it live will be 10.26180/14073236)

## References and Notes

1. M. Pavan, D. Bassani, M. Sturlese, S. Moro, From the Wuhan-Hu-1 strain to the XD and XE variants: is targeting the SARS-CoV-2 spike protein still a pharmaceutically relevant option against COVID-19? J Enzyme Inhib Med Chem 37, 1704–1714 (2022).

2. M. Gavriatopoulou et al., Emerging treatment strategies for COVID-19 infection. Clin Exp Med 21, 167–179 (2021).

3. J. Majumder, T. Minko, Recent Developments on Therapeutic and Diagnostic Approaches for COVID-19. Aaps j 23, 14 (2021).

4. N. Zhu et al., A Novel Coronavirus from Patients with Pneumonia in China, 2019. N Engl J Med 382, 727–733 (2020).

5. T. M. Delorey et al., COVID-19 tissue atlases reveal SARS-CoV-2 pathology and cellular targets. Nature 595, 107–113 (2021).

6. J. C. Melms et al., A molecular single-cell lung atlas of lethal COVID-19. Nature 595, 114–119 (2021).

7. A. Calkovska, M. Kolomaznik, V. Calkovsky, Alveolar type II cells and pulmonary surfactant in COVID-19 era. Physiol Res 70, S195–s208 (2021).

8. L. Guillot et al., Alveolar epithelial cells: master regulators of lung homeostasis. Int J Biochem Cell Biol 45, 2568–2573 (2013).

9. P. Smith, D. Heath, H. Moosavi, The Clara cell. Thorax 29, 147–163 (1974).

10. A. Hamvas et al., Developmental and genetic regulation of human surfactant protein B in vivo. Neonatology 95, 117–124 (2009).

11. S. Guttentag, Posttranslational regulation of surfactant protein B expression. Semin Perinatol 32, 367–370 (2008).

12. S. Reuter, C. Moser, M. Baack, Respiratory distress in the newborn. Pediatr Rev 35, 417–428; quiz 429 (2014).

13. V. Kavvadia, A. Greenough, Y. Itakura, G. Dimitriou, Neonatal lung function in very immature infants with and without RDS. J Perinat Med 27, 382–387 (1999).

14. J. P. de Winter, I. T. Merth, R. Brand, P. H. Quanjer, Functional residual capacity and static compliance during the first year in preterm infants treated with surfactant. Am J Perinatol 17, 377–384 (2000).

15. J. Johansson, T. Curstedt, Synthetic surfactants with SP-B and SP-C analogues to enable worldwide treatment of neonatal respiratory distress syndrome and other lung diseases. J Intern Med 285, 165–186 (2019).

16. P. Schousboe et al., Assessment of pulmonary surfactant in COVID-19 patients. Crit Care 24, 552 (2020).

17. W. J. Davidson, et al., Exogenous pulmonary surfactant for the treatment of adult patients with acute respiratory distress syndrome: results of a meta-analysis. *Crit Care* 10, R41 (2006).

18. A. Ghati et al., Exogenous pulmonary surfactant: A review focused on adjunctive therapy for severe acute respiratory syndrome coronavirus 2 including SP-A and SP-D as added clinical marker. Curr Opin Colloid Interface Sci 51, 101413 (2021).

19. S. L. Leibel, R. N. McVicar, A. M. Winquist, W. D. Niles, E. Y. Snyder, Generation of Complete Multi-Cell Type Lung Organoids From Human Embryonic and Patient-Specific Induced Pluripotent Stem Cells for Infectious Disease Modeling and Therapeutics Validation. Curr Protoc Stem Cell Biol 54, e118 (2020).

20. S. L. Leibel et al., Reversal of Surfactant Protein B Deficiency in Patient Specific Human Induced Pluripotent Stem Cell Derived Lung Organoids by Gene Therapy. Sci Rep 9, 13450 (2019).

21. Y. Yamamoto et al., Long-term expansion of alveolar stem cells derived from human iPS cells in organoids. Nat Methods 14, 1097–1106 (2017).

22. K. B. McCauley, F. Hawkins, D. N. Kotton, Derivation of Epithelial-Only Airway Organoids from Human Pluripotent Stem Cells. Curr Protoc Stem Cell Biol 45, e51 (2018).

23. J. K. Fiege et al., Single cell resolution of SARS-CoV-2 tropism, antiviral responses, and susceptibility to therapies in primary human airway epithelium. PLoS Pathog 17, e1009292 (2021).

24. R. J. Mason, Pathogenesis of COVID-19 from a cell biology perspective. Eur Respir J 55, (2020).

25. R. L. Chua et al., COVID-19 severity correlates with airway epithelium-immune cell interactions identified by single-cell analysis. Nat Biotechnol 38, 970–979 (2020).

26. N. G. Ravindra et al., Single-cell longitudinal analysis of SARS-CoV-2 infection in human airway epithelium identifies target cells, alterations in gene expression, and cell state changes. PLoS Biol 19, e3001143 (2021).

27. L. Milross et al., Post-mortem lung tissue: the fossil record of the pathophysiology and immunopathology of severe COVID-19. The Lancet Respiratory Medicine 10, 95–106 (2022).

28. T. Mauad et al., Tracking the time course of pathological patterns of lung injury in severe COVID-19. Respir Res 22, 32 (2021).

29. V. Cagno, SARS-CoV-2 cellular tropism. The Lancet Microbe 1, e2–e3 (2020).

30. S. Yang et al., Transcriptomic analysis reveals novel mechanisms of SARS-CoV-2 infection in human lung cells. Immunity, Inflammation and Disease 8, 753–762 (2020).

31. A. Kubo et al., Development of definitive endoderm from embryonic stem cells in culture. Development 131, 1651–1662 (2004).

32. M. D. Green et al., Generation of anterior foregut endoderm from human embryonic and induced pluripotent stem cells. Nat Biotechnol 29, 267–272 (2011).

33. S. A. Rankin et al., A Retinoic Acid-Hedgehog Cascade Coordinates Mesoderm-Inducing Signals and Endoderm Competence during Lung Specification. Cell Rep 16, 66–78 (2016).

34. S. Gotoh et al., Generation of alveolar epithelial spheroids via isolated progenitor cells from human pluripotent stem cells. Stem Cell Reports 3, 394–403 (2014).

35. S. L. Leibel, R. N. McVicar, A. M. Winquist, E. Y. Snyder, Generation of 3D Whole Lung Organoids from Induced Pluripotent Stem Cells for Modeling Lung Developmental Biology and Disease. J Vis Exp, (2021).

36. P. Minoo et al., Physical and functional interactions between homeodomain NKX2.1 and winged helix/forkhead FOXA1 in lung epithelial cells. Mol Cell Biol 27, 2155–2165 (2007).

37. K. J. Travaglini et al., A molecular cell atlas of the human lung from single-cell RNA sequencing. Nature 587, 619–625 (2020).

38. K. Hurley et al., Reconstructed Single-Cell Fate Trajectories Define Lineage Plasticity Windows during Differentiation of Human PSC-Derived Distal Lung Progenitors. Cell Stem Cell 26, 593–608.e598 (2020).

39. M. E. Ardini-Poleske et al., LungMAP: The Molecular Atlas of Lung Development Program. Am J Physiol Lung Cell Mol Physiol 313, L733–l740 (2017).

40. Y. Du, M. Guo, J. A. Whitsett, Y. Xu, ’LungGENS’: a web-based tool for mapping single-cell gene expression in the developing lung. Thorax 70, 1092–1094 (2015).

41. P. A. Reyfman et al., Single-Cell Transcriptomic Analysis of Human Lung Provides Insights into the Pathobiology of Pulmonary Fibrosis. American Journal of Respiratory and Critical Care Medicine 199, 1517–1536 (2019).

42. H. B. Schiller et al., The Human Lung Cell Atlas: A High-Resolution Reference Map of the Human Lung in Health and Disease. American Journal of Respiratory Cell and Molecular Biology 61, 31–41 (2019).

43. Y. Xu, et al., Single-cell RNA sequencing identifies diverse roles of epithelial cells in idiopathic pulmonary fibrosis. JCI Insight 1, (2017).

44. K. B. McCauley et al., Single-Cell Transcriptomic Profiling of Pluripotent Stem Cell-Derived SCGB3A2+ Airway Epithelium. Stem Cell Reports 10, 1579–1595 (2018).

45. Y. Zhou et al., Metascape provides a biologist-oriented resource for the analysis of systems-level datasets. Nat Commun 10, 1523 (2019).

46. L. Riva et al., Discovery of SARS-CoV-2 antiviral drugs through large-scale compound repurposing. Nature 586, 113–119 (2020).

47. G. Nunnari et al., Network perturbation analysis in human bronchial epithelial cells following SARS-CoV2 infection. Exp Cell Res 395, 112204 (2020).

48. C. Ota et al., Dynamic expression of HOPX in alveolar epithelial cells reflects injury and repair during the progression of pulmonary fibrosis. Scientific Reports 8, 12983 (2018).

49. Z. Zhao et al., Single-cell analysis identified lung progenitor cells in COVID-19 patients. Cell Prolif 53, e12931 (2020).

50. K. Dunigan et al., The thioredoxin reductase inhibitor auranofin induces heme oxygenase-1 in lung epithelial cells via Nrf2-dependent mechanisms. Am J Physiol Lung Cell Mol Physiol 315, L545–l552 (2018).

51. J. D. Chandler et al., Metabolic pathways of lung inflammation revealed by high-resolution metabolomics (HRM) of H1N1 influenza virus infection in mice. Am J Physiol Regul Integr Comp Physiol 311, R906–r916 (2016).

52. T. Ma et al., RNA-Seq Analysis of Influenza A Virus-Induced Transcriptional Changes in Mice Lung and Its Possible Implications for the Virus Pathogenicity in Mice. Viruses 13, 2031 (2021).

53. H. Blasco et al., The specific metabolome profiling of patients infected by SARS-COV-2 supports the key role of tryptophan-nicotinamide pathway and cytosine metabolism. Sci Rep 10, 16824 (2020).

54. Q. Feng, L. Li, X. Wang, Identifying Pathways and Networks Associated With the SARS-CoV-2 Cell Receptor ACE2 Based on Gene Expression Profiles in Normal and SARS-CoV-2-Infected Human Tissues. Front Mol Biosci 7, 568954 (2020).

55. Y. Lissanu Deribe et al., Mutations in the SWI/SNF complex induce a targetable dependence on oxidative phosphorylation in lung cancer. Nature Medicine 24, 1047–1057 (2018).

56. M. Hu et al., Respiratory syncytial virus co-opts host mitochondrial function to favour infectious virus production. eLife 8, e42448 (2019).

57. R. Cecchini, A. L. Cecchini, SARS-CoV-2 infection pathogenesis is related to oxidative stress as a response to aggression. Med Hypotheses 143, 110102 (2020).

58. M. Weitnauer, V. Mijošek, A. H. Dalpke, Control of local immunity by airway epithelial cells. Mucosal Immunology 9, 287–298 (2016).

59. R. J. Hewitt, C. M. Lloyd, Regulation of immune responses by the airway epithelial cell landscape. Nature Reviews Immunology 21, 347–362 (2021).

60. K. Brune, J. Frank, A. Schwingshackl, J. Finigan, V. K. Sidhaye, Pulmonary epithelial barrier function: some new players and mechanisms. Am J Physiol Lung Cell Mol Physiol 308, L731–745 (2015).

61. S. Satarker, M. Nampoothiri, Structural Proteins in Severe Acute Respiratory Syndrome Coronavirus-2. Arch Med Res 51, 482–491 (2020).

62. N. Redondo, S. Zaldívar-López, J. J. Garrido, M. Montoya, SARS-CoV-2 Accessory Proteins in Viral Pathogenesis: Knowns and Unknowns. Front Immunol 12, 708264 (2021).

63. D. Kim et al., A high-resolution temporal atlas of the SARS-CoV-2 translatome and transcriptome. Nature Communications 12, 5120 (2021).

64. T. M. Clausen et al., SARS-CoV-2 Infection Depends on Cellular Heparan Sulfate and ACE2. bioRxiv, (2020).

65. M. Hoffmann et al., SARS-CoV-2 Cell Entry Depends on ACE2 and TMPRSS2 and Is Blocked by a Clinically Proven Protease Inhibitor. Cell 181, 271–280.e278 (2020).

66. H. P. Jia et al., ACE2 Receptor Expression and Severe Acute Respiratory Syndrome Coronavirus Infection Depend on Differentiation of Human Airway Epithelia. Journal of Virology 79, 14614–14621 (2005).

67. J. Lan et al., Structure of the SARS-CoV-2 spike receptor-binding domain bound to the ACE2 receptor. Nature 581, 215–220 (2020).

68. W. Li et al., Angiotensin-converting enzyme 2 is a functional receptor for the SARS coronavirus. Nature 426, 450–454 (2003).

69. R. Yan et al., Structural basis for the recognition of SARS-CoV-2 by full-length human ACE2. Science 367, 1444–1448 (2020).

70. Z. Zhang et al., Clinical analysis and pluripotent stem cells-based model reveal possible impacts of ACE2 and lung progenitor cells on infants vulnerable to COVID-19. Theranostics 11, 2170–2181 (2021).

71. Y. Zhao et al., Single-cell RNA expression profiling of ACE2, the putative receptor of Wuhan 2019-nCov. bioRxiv, 2020.2001.2026.919985 (2020).

72. C. G. K. Ziegler et al., SARS-CoV-2 receptor ACE2 is an interferon-stimulated gene in human airway epithelial cells and is detected in specific cell subsets across tissues. Cell.

73. C. B. Jackson, M. Farzan, B. Chen, H. Choe, Mechanisms of SARS-CoV-2 entry into cells. Nature Reviews Molecular Cell Biology 23, 3–20 (2022).

74. M. Hoffmann, H. Kleine-Weber, S. Pöhlmann, A Multibasic Cleavage Site in the Spike Protein of SARS-CoV-2 Is Essential for Infection of Human Lung Cells. Mol Cell 78, 779–784.e775 (2020).

75. J. Shang et al., Cell entry mechanisms of SARS-CoV-2. Proc Natl Acad Sci U S A 117, 11727–11734 (2020).

76. C. Yang et al., Kidney injury molecule-1 is a potential receptor for SARS-CoV-2. Journal of Molecular Cell Biology 13, 185–196 (2021).

77. S. Wang et al., AXL is a candidate receptor for SARS-CoV-2 that promotes infection of pulmonary and bronchial epithelial cells. Cell Res 31, 126–140 (2021).

78. L. Cantuti-Castelvetri et al., Neuropilin-1 facilitates SARS-CoV-2 cell entry and infectivity. Science 370, 856–860 (2020).

79. J. L. Daly et al., Neuropilin-1 is a host factor for SARS-CoV-2 infection. Science 370, 861–865 (2020).

80. Y. Zhang, R. Yan, Q. Zhou, ACE2, B0AT1, and SARS-CoV-2 spike protein: Structural and functional implications. Current Opinion in Structural Biology 74, 102388 (2022).

81. K. Prasad, S. Y. AlOmar, E. A. Almuqri, H. A. Rudayni, V. Kumar, Genomics-guided identification of potential modulators of SARS-CoV-2 entry proteases, TMPRSS2 and Cathepsins B/L. PLoS One 16, e0256141 (2021).

82. M.-M. Zhao et al., Cathepsin L plays a key role in SARS-CoV-2 infection in humans and humanized mice and is a promising target for new drug development. Signal Transduction and Targeted Therapy 6, 134 (2021).

83. B. Cao, L. Zhang, H. Liu, S. Ma, K. Mi, The Dynamic Expression of Potential Mediators of Severe Acute Respiratory Syndrome Coronavirus 2 Cellular Entry in Fetal, Neonatal, and Adult Rhesus Monkeys. Front Genet 11, 607479 (2020).

84. Z. Chen et al., Function of HAb18G/CD147 in invasion of host cells by severe acute respiratory syndrome coronavirus. J Infect Dis 191, 755–760 (2005).

85. K. Wang et al., CD147-spike protein is a novel route for SARS-CoV-2 infection to host cells. Signal Transduct Target Ther 5, 283 (2020).

86. S. Gayle et al., Identification of apilimod as a first-in-class PIKfyve kinase inhibitor for treatment of B-cell non-Hodgkin lymphoma. Blood 129, 1768–1778 (2017).

87. P. T. Huang, S. Einav, C. R. M. Asquith, PIKfyve: a lipid kinase target for COVID-19, cancer and neurodegenerative disorders. Nat Rev Drug Discov 20, 730 (2021).

88. S. Krishna et al., PIKfyve Regulates Vacuole Maturation and Nutrient Recovery following Engulfment. Dev Cell 38, 536–547 (2016).

89. T. Duong le, A. T. Leung, B. Langdahl, Cathepsin K Inhibition: A New Mechanism for the Treatment of Osteoporosis. Calcif Tissue Int 98, 381–397 (2016).

90. J. H. Beigel et al., Remdesivir for the Treatment of Covid-19 — Final Report. New England Journal of Medicine 383, 1813–1826 (2020).

91. L. Riva, et al., A Large-scale Drug Repositioning Survey for SARS-CoV-2 Antivirals. bioRxiv, 2020.2004.2016.044016 (2020).

92. J. Cubuk et al., The SARS-CoV-2 nucleocapsid protein is dynamic, disordered, and phase separates with RNA. Nature Communications 12, 1936 (2021).

93. B. Boson et al., The SARS-CoV-2 envelope and membrane proteins modulate maturation and retention of the spike protein, allowing assembly of virus-like particles. Journal of Biological Chemistry 296, 100111 (2021).

94. H. Geng et al., SARS-CoV-2 ORF8 Forms Intracellular Aggregates and Inhibits IFNγ-Induced Antiviral Gene Expression in Human Lung Epithelial Cells. Front Immunol 12, 679482 (2021).

95. U. Timilsina, S. Umthong, E. B. Ivey, B. Waxman, S. Stavrou, SARS-CoV-2 ORF7a potently inhibits the antiviral effect of the host factor SERINC5. Nature Communications 13, 2935 (2022).

96. I. Rusinova et al., Interferome v2.0: an updated database of annotated interferon-regulated genes. Nucleic Acids Res 41, D1040–1046 (2013).

97. P. Hertzog, S. Forster, S. Samarajiwa, Systems biology of interferon responses. J Interferon Cytokine Res 31, 5–11 (2011).

98. A. Krämer, J. Green, J. Pollard, Jr., S. Tugendreich, Causal analysis approaches in Ingenuity Pathway Analysis. Bioinformatics 30, 523–530 (2014).

99. A. Bass, Y. Liu, S. Dakshanamurthy, Single-Cell and Bulk RNASeq Profiling of COVID-19 Patients Reveal Immune and Inflammatory Mechanisms of Infection-Induced Organ Damage. Viruses 13, (2021).

100. S. Li et al., SARS-CoV-2 triggers inflammatory responses and cell death through caspase-8 activation. Signal Transduction and Targeted Therapy 5, 235 (2020).

101. B. Sposito et al., The interferon landscape along the respiratory tract impacts the severity of COVID-19. Cell 184, 4953–4968.e4916 (2021).

102. N. Sever, G. Miličić, N. O. Bodnar, X. Wu, T. A. Rapoport, Mechanism of Lamellar Body Formation by Lung Surfactant Protein B. Mol Cell 81, 49–66.e48 (2021).

103. K. Osanai, S. Mizuno, H. Toga, K. Takahashi, Trafficking of newly synthesized surfactant protein B to the lamellar body in alveolar type II cells. Cell Tissue Res 381, 427–438 (2020).

104. C. D. Foster, P. X. Zhang, L. W. Gonzales, S. H. Guttentag, In vitro surfactant protein B deficiency inhibits lamellar body formation. Am J Respir Cell Mol Biol 29, 259–266 (2003).

105. A. L. Johnson et al., Post-Translational Processing of Surfactant Protein-C Proprotein. American Journal of Respiratory Cell and Molecular Biology 24, 253–263 (2001).

106. A. Hamvas, L. M. Nogee, D. E. deMello, F. S. Cole, Pathophysiology and treatment of surfactant protein-B deficiency. Biol Neonate 67 Suppl 1, 18–31 (1995).

107. S. H. Guttentag, M. F. Beers, B. M. Bieler, P. L. Ballard, Surfactant protein B processing in human fetal lung. Am J Physiol 275, L559–566 (1998).

108. J. M. Coya et al., Natural Anti-Infective Pulmonary Proteins: In Vivo Cooperative Action of Surfactant Protein SP-A and the Lung Antimicrobial Peptide SP-B^N^. The Journal of Immunology 195, 1628–1636 (2015).

109. M. Ikegami, J. A. Whitsett, P. C. Martis, T. E. Weaver, Reversibility of lung inflammation caused by SP-B deficiency. Am J Physiol Lung Cell Mol Physiol 289, L962–970 (2005).

110. L. Yang et al., Surfactant protein B propeptide contains a saposin-like protein domain with antimicrobial activity at low pH. J Immunol 184, 975–983 (2010).

111. L. M. Nogee et al., A mutation in the surfactant protein B gene responsible for fatal neonatal respiratory disease in multiple kindreds. J Clin Invest 93, 1860–1863 (1994).

112. L. M. Nogee, S. E. Wert, S. A. Proffit, W. M. Hull, J. A. Whitsett, Allelic heterogeneity in hereditary surfactant protein B (SP-B) deficiency. Am J Respir Crit Care Med 161, 973–981 (2000).

113. A. Jacob et al., Differentiation of Human Pluripotent Stem Cells into Functional Lung Alveolar Epithelial Cells. Cell Stem Cell 21, 472–488.e410 (2017).

114. L. Y. Drake, H. Kita, IL-33: biological properties, functions, and roles in airway disease. Immunol Rev 278, 173–184 (2017).

115. S. J. Gurczynski, B. B. Moore, IL-17 in the lung: the good, the bad, and the ugly. Am J Physiol Lung Cell Mol Physiol 314, L6–l16 (2018).

116. C. Hoffman et al., Interleukin-19: a constituent of the regulome that controls antigen presenting cells in the lungs and airway responses to microbial products. PLoS One 6, e27629 (2011).

117. B. A. Khalil, N. M. Elemam, A. A. Maghazachi, Chemokines and chemokine receptors during COVID-19 infection. Computational and Structural Biotechnology Journal 19, 976–988 (2021).

118. S. J. Attarian et al., Mutations in the thyroid transcription factor gene NKX2-1 result in decreased expression of SFTPB and SFTPC. Pediatr Res 84, 419–425 (2018).

119. J. B. Tagne et al., Genome-wide analyses of Nkx2-1 binding to transcriptional target genes uncover novel regulatory patterns conserved in lung development and tumors. PLoS One 7, e29907 (2012).

120. Z. Inde et al., Age-dependent regulation of SARS-CoV-2 cell entry genes and cell death programs correlates with COVID-19 severity. Science Advances 7, eabf8609 (2021).

121. J. M. Emeny, M. J. Morgan, Regulation of the interferon system: evidence that Vero cells have a genetic defect in interferon production. J Gen Virol 43, 247–252 (1979).

122. R. Ramanathan, J. J. Bhatia, K. Sekar, F. R. Ernst, Mortality in preterm infants with respiratory distress syndrome treated with poractant alfa, calfactant or beractant: a retrospective study. J Perinatol 33, 119–125 (2013).

123. M. Farahani et al., Molecular pathways involved in COVID-19 and potential pathway-based therapeutic targets. Biomed Pharmacother 145, 112420 (2022).

124. S. Montazersaheb et al., COVID-19 infection: an overview on cytokine storm and related interventions. Virology Journal 19, 92 (2022).

125. J. R. Wright, Pulmonary surfactant: a front line of lung host defense. J Clin Invest 111, 1453–1455 (2003).

126. J. R. Wright, Immunomodulatory functions of surfactant. Physiol Rev 77, 931–962 (1997).

127. M. Numata, H. W. Chu, A. Dakhama, D. R. Voelker, Pulmonary surfactant phosphatidylglycerol inhibits respiratory syncytial virus-induced inflammation and infection. Proc Natl Acad Sci U S A 107, 320–325 (2010).

128. R. G. Spragg et al., Effect of recombinant surfactant protein C-based surfactant on the acute respiratory distress syndrome. N Engl J Med 351, 884–892 (2004).

129. D. R. Voelker, M. Numata, Phospholipid regulation of innate immunity and respiratory viral infection. J Biol Chem 294, 4282–4289 (2019).

130. U. Mirastschijski, R. Dembinski, K. Maedler, Lung Surfactant for Pulmonary Barrier Restoration in Patients With COVID-19 Pneumonia. Front Med (Lausanne) 7, 254 (2020).

131. S. Wang et al., The Role of Pulmonary Surfactants in the Treatment of Acute Respiratory Distress Syndrome in COVID-19. Front Pharmacol 12, 698905 (2021).

132. K. Pramod et al., Surfactant-based prophylaxis and therapy against COVID-19: A possibility. Med Hypotheses 143, 110081–110081 (2020).

133. Z. G. Dessie, T. Zewotir, Mortality-related risk factors of COVID-19: a systematic review and meta-analysis of 42 studies and 423,117 patients. BMC Infect Dis 21, 855 (2021).

134. M. Webb Hooper, A. M. Nápoles, E. J. Pérez-Stable, COVID-19 and Racial/Ethnic Disparities. Jama 323, 2466–2467 (2020).

135. R. T. Gandhi, P. N. Malani, C. Del Rio, COVID-19 Therapeutics for Nonhospitalized Patients. Jama 327, 617–618 (2022).

136. A. Daher et al., Clinical course of COVID-19 patients needing supplemental oxygen outside the intensive care unit. Sci Rep 11, 2256 (2021).

137. L. A. Hajjar et al., Intensive care management of patients with COVID-19: a practical approach. Ann Intensive Care 11, 36 (2021).

138. E. Takashita et al., Efficacy of Antibodies and Antiviral Drugs against Covid-19 Omicron Variant. N Engl J Med 386, 995–998 (2022).

139. U. Radzikowska et al., Distribution of ACE2, CD147, CD26, and other SARS-CoV-2 associated molecules in tissues and immune cells in health and in asthma, COPD, obesity, hypertension, and COVID-19 risk factors. Allergy 75, 2829–2845 (2020).

140. A. Wang et al., Single-cell multiomic profiling of human lungs reveals cell-type-specific and age-dynamic control of SARS-CoV2 host genes. Elife 9, (2020).

141. A. Purkayastha et al., Direct Exposure to SARS-CoV-2 and Cigarette Smoke Increases Infection Severity and Alters the Stem Cell-Derived Airway Repair Response. Cell Stem Cell 27, 869–875.e864 (2020).

142. J. C. Smith et al., Cigarette Smoke Exposure and Inflammatory Signaling Increase the Expression of the SARS-CoV-2 Receptor ACE2 in the Respiratory Tract. Developmental Cell 53, 514–529.e513 (2020).

143. J. C. Smith, J. M. Sheltzer, Cigarette smoke triggers the expansion of a subpopulation of respiratory epithelial cells that express the SARS-CoV-2 receptor ACE2. bioRxiv, 2020.2003.2028.013672 (2020).

144. J. J. Baczenas et al., Propagation of SARS-CoV-2 in Calu-3 Cells to Eliminate Mutations in the Furin Cleavage Site of Spike. Viruses 13, (2021).

145. S. G. P. Funnell et al., A cautionary perspective regarding the isolation and serial propagation of SARS-CoV-2 in Vero cells. NPJ Vaccines 6, 83 (2021).

146. S. Berg et al., ilastik: interactive machine learning for (bio)image analysis. Nat Methods 16, 1226–1232 (2019).

147. R. Haase et al., CLIJ: GPU-accelerated image processing for everyone. Nat Methods 17, 5–6 (2020).

148. A. Subramanian et al., Gene set enrichment analysis: A knowledge-based approach for interpreting genome-wide expression profiles. Proceedings of the National Academy of Sciences 102, 15545–15550 (2005).

149. T. Stuart et al., Comprehensive Integration of Single-Cell Data. Cell 177, 1888–1902 e1821 (2019).

150. I. Korsunsky et al., Fast, sensitive and accurate integration of single-cell data with Harmony. Nat Methods 16, 1289–1296 (2019).

151. M. G. Schimek et al., TopKLists: a comprehensive R package for statistical inference, stochastic aggregation, and visualization of multiple omics ranked lists. Stat Appl Genet Mol Biol 14, 311–316 (2015).

152. C. Commisso, R. J. Flinn, D. Bar-Sagi, Determining the macropinocytic index of cells through a quantitative image-based assay. Nat Protoc 9, 182–192 (2014).

153. K. M. O. Galenkamp, B. Alas, C. Commisso, Quantitation of Macropinocytosis in Cancer Cells. Methods Mol Biol 1928, 113–123 (2019).

154. K. M. O. Galenkamp, C. M. Galapate, Y. Zhang, C. Commisso, Automated Imaging and Analysis for the Quantification of Fluorescently Labeled Macropinosomes. J Vis Exp, (2021).

