## Supplemental Figures and Tables for "The lung employs an intrinsic surfactant-mediated inflammatory response for viral defense"

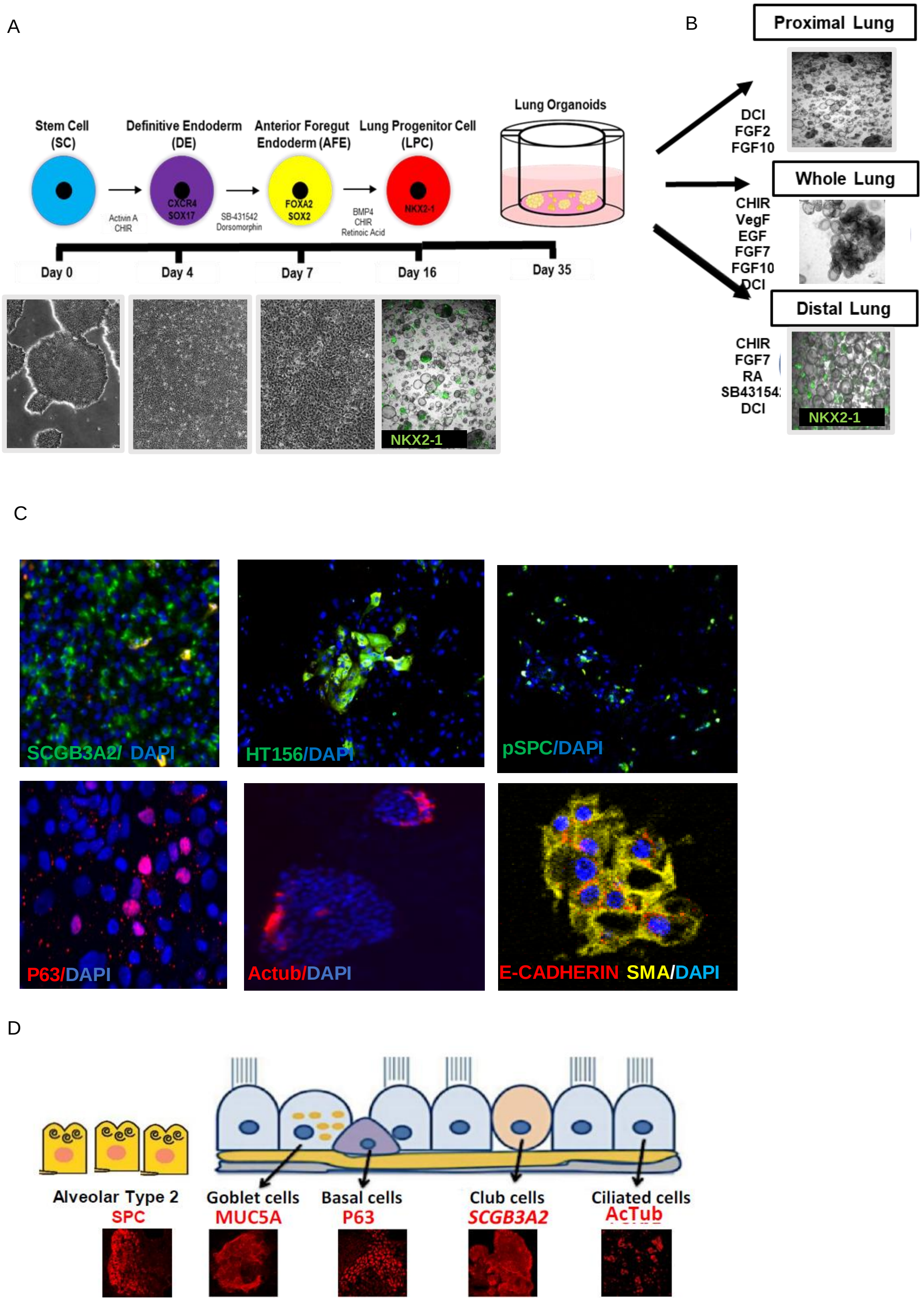

**FIGURE S1: Differentiation of human pluripotent stem cells (hPSC) into multi-cell type 3D lung organoids (LOs).** (A) Schematic for directed differentiation of hPSCs into 3D LOs with corresponding phase contrast images of each step, including undifferentiated hPSCs, definitive endoderm, anterior foregut endoderm, and lung progenitor cells (LPC). Image of 3D LPCs shows *NKX2-1* via GFP expression. (B) Live cell phase contrast images of the representative morphology of 3 different 3D lung organoid types: proximal (PLO), distal (DLO) and whole lung organoids (WLO). (C) Immunostaining of dissociated 3D WLOs with some representative lung epithelial and mesenchymal markers. Their respective cell types are shown in the schematic in (D). (D) Schematic of the epithelial cells in the PLOs and DLOs with representative immunofluorescent images of the epithelial markers found in the DLOs (alveolar type 2 cells) and PLOs (airway cells).

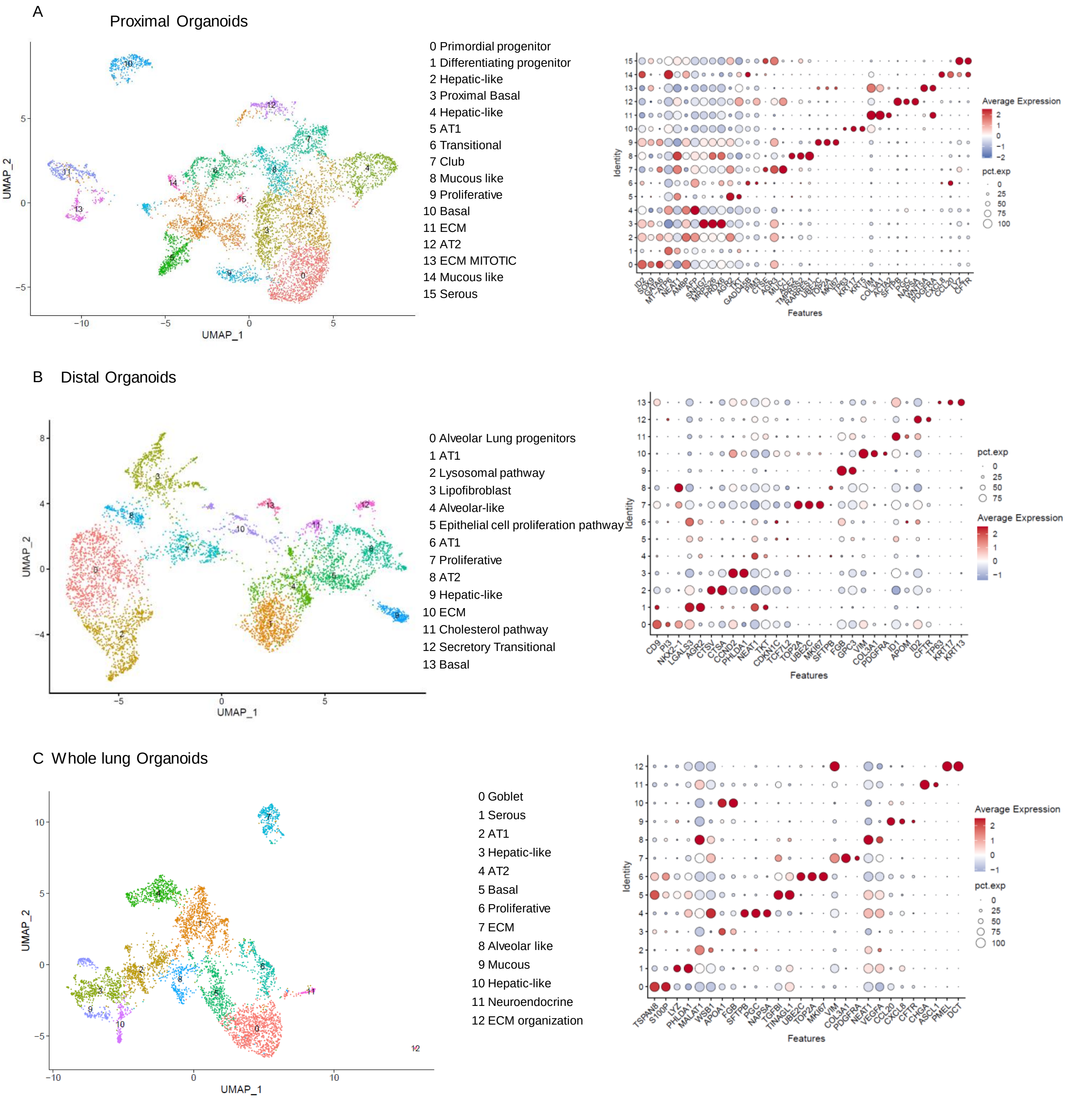

**FIGURE S2: Transcriptional characterization of LOs.** (A) Uniform manifold approximation and projection (UMAP) of hPSC-PLOs and characteristic genes per cluster. (B) UMAP of hPSC-DLOs and characteristic genes per cluster. (C) UMAP of hPSC-WLOs and characteristic genes per cluster.

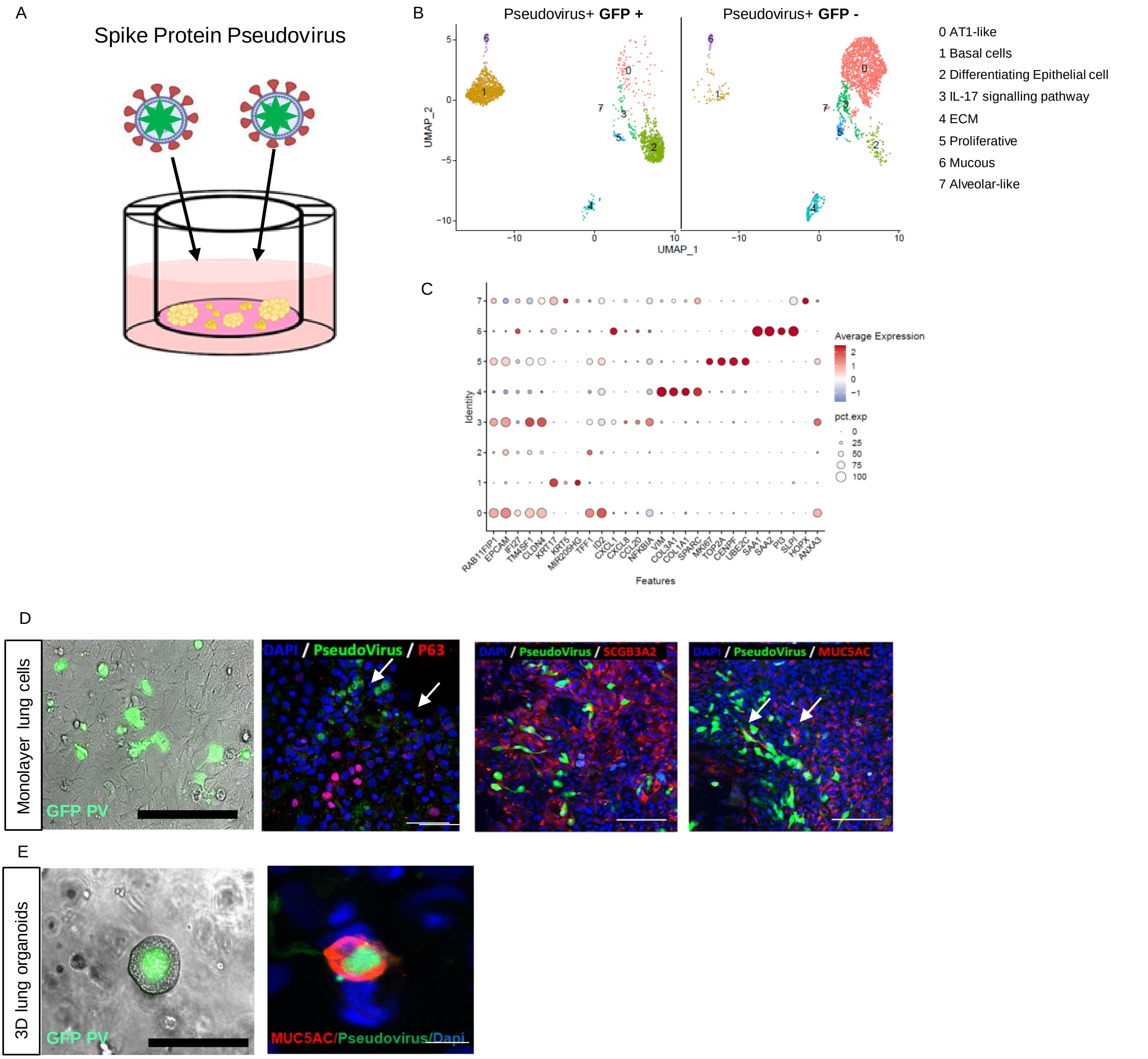

**FIGURE S3: Infection of hPSC-derived lung organoids (LOs) with SARS-CoV-2 pseudovirus.** (A) Schematic for SARS-CoV-2 pseudovirus infection in human lung organoids (LO). LOs were cultured in serum free basal medium and infected with SARS-CoV-2 pseudovirus for 24 hours. (B) Uniform manifold approximation and projection (UMAP) of 3D hPSC-PLOs infected with pseudovirus conjugated to a GFP reporter. The clusters show the tropism of the SARS-CoV-2 spike protein. The UMAP cluster on the left represents LOs that were exposed to pseudovirus and infected. The UMAP cluster on the right represents LOs that were exposed to the pseudovirus but were not infected. (C) Characteristic genes identifying each cluster. (D) Dual immunostaining of 3D PLOs (acutely dissociated to enable visualization of individual cells in monolayer without extensive overlapping of cells) for both identifying lung epithelial markers and pseudovirus-GFP. (E) Representative immunostaining of live, intact 3D PLOs with goblet cell marker MUC5AC and pseudovirus-GFP.

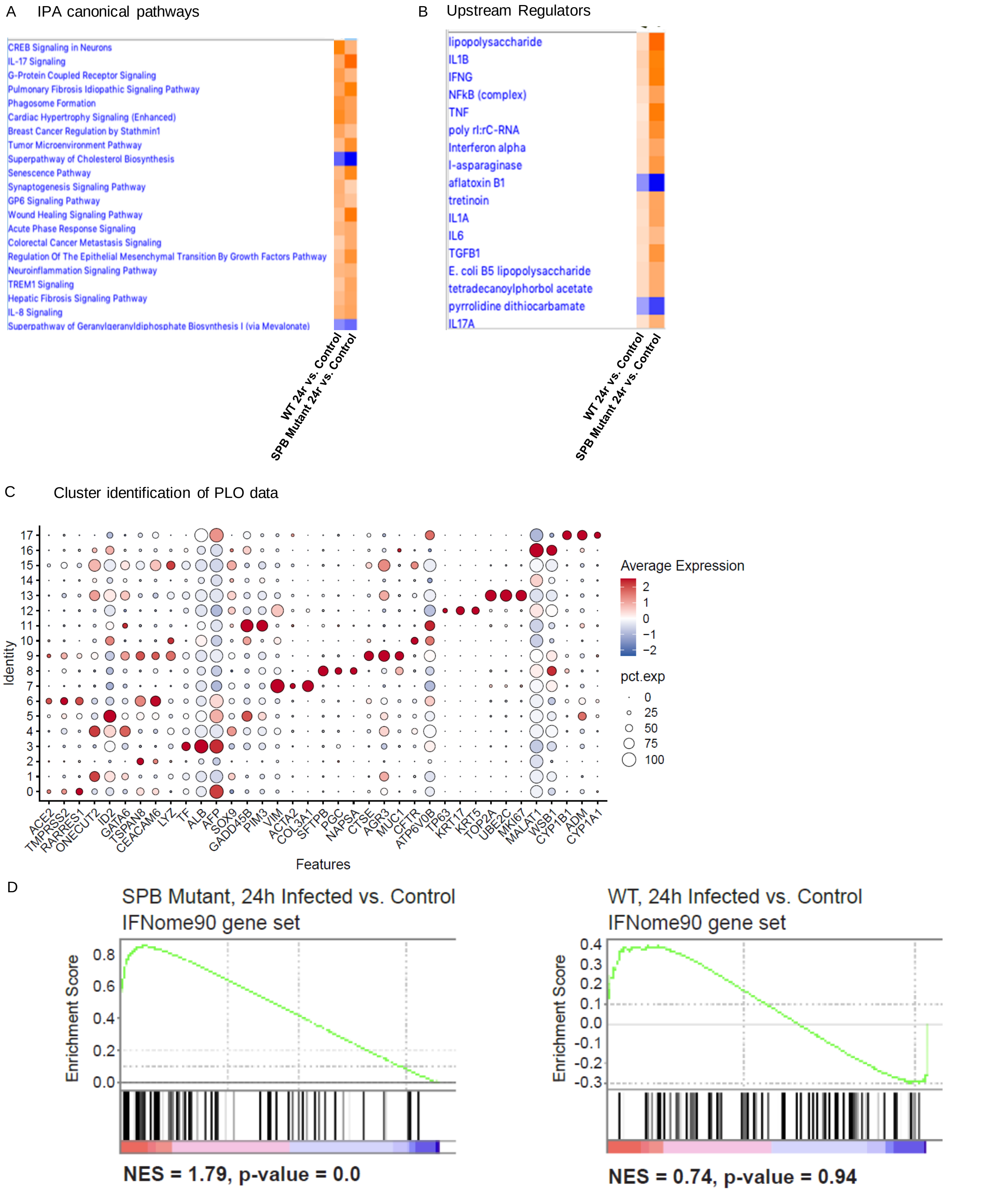

**FIGURE S4: Gene enrichment pathways of infected WT and surfactant protein B (SP-B)-deficient LO cells (by virtue of a deletion mutation in a patient) vs mock infected controls.** (A) Ingenuity pathway analysis (IPA) canonical pathways comparing the gene expression patterns of WT vs. SP-B deficient PLOs 24 hours post infection (hpi) with a replication-competent SARS-CoV-2 virus vs. mock-infected controls (B) IPA upstream regulators of the 24 hpi and mock infected wt and SP-B deficient PLOs. (C) Characteristic genes per cluster for the PLO single cell RNA sequencing data. (D) IFNome90 gene set enrichment scores of 24 hpi vs mock infected wild type and SP-B deficient PLOs. Positive NES defines interferon enrichment.

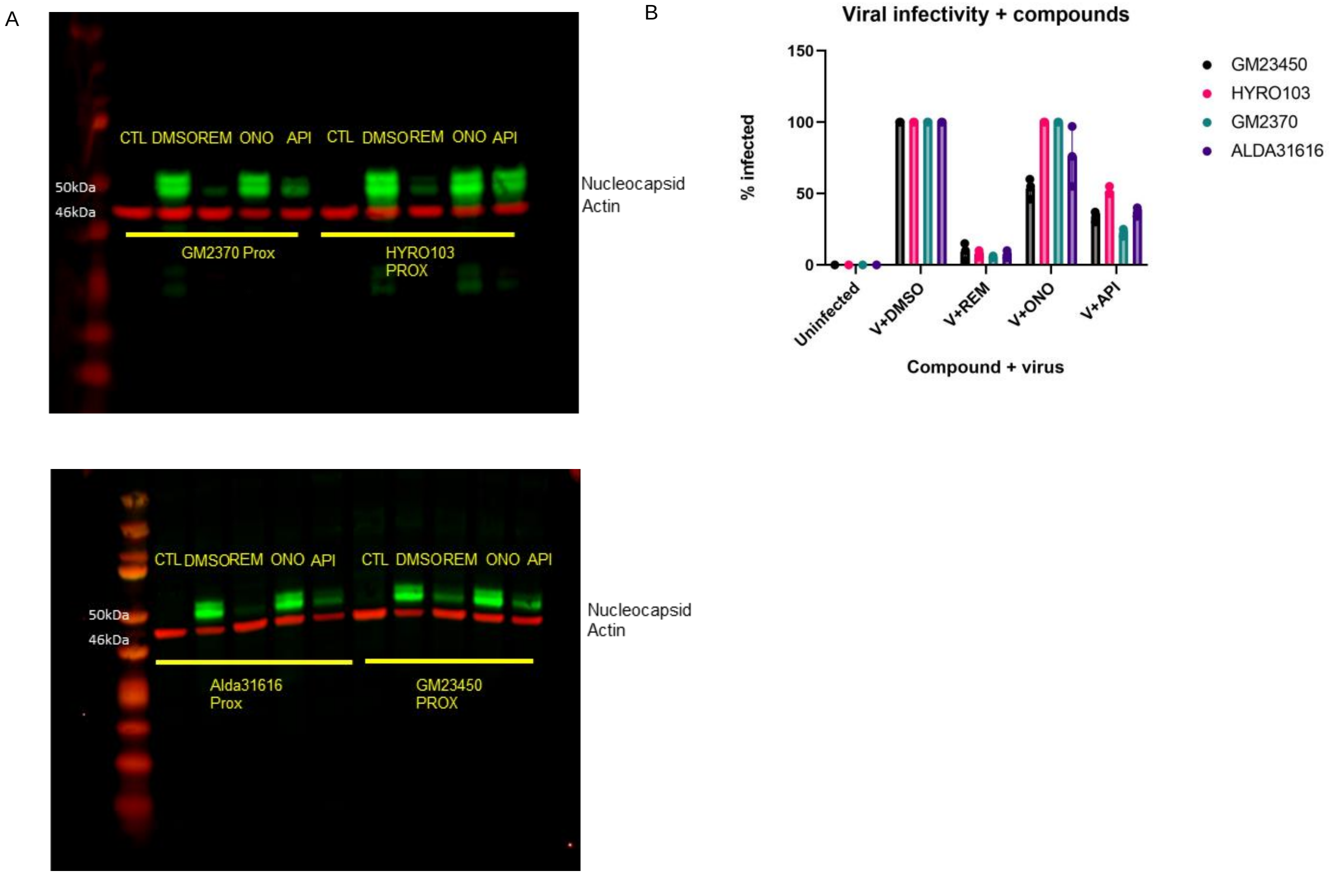

**FIGURE S5: Compounds blocking various routes of viral entry and processing show different efficacies in reducing viral infections.** (A) Western blot of PLOs from different racial groups, pre-treated with the endocytic blocking agent (apilimod [“API”]), the cathepsin B blocking agent (ONO5334 [“ONO”]), or an agent that blocks viral replication (Remdesivir [“REM”]) 24 hpi with SARS-CoV-2 (“V”). (B) FACS results of infected cells after exposure to the same compounds. The results are normalized to virus + DMSO. The data are representative of three independent experiments. PLO GM23450 was derived from hiPSCs generated from an African-American female, HYRO103 from a Hispanic male, GM2370 from a Caucasian female, and ALDA31616 from a Caucasian male (Supplementary Table 1.

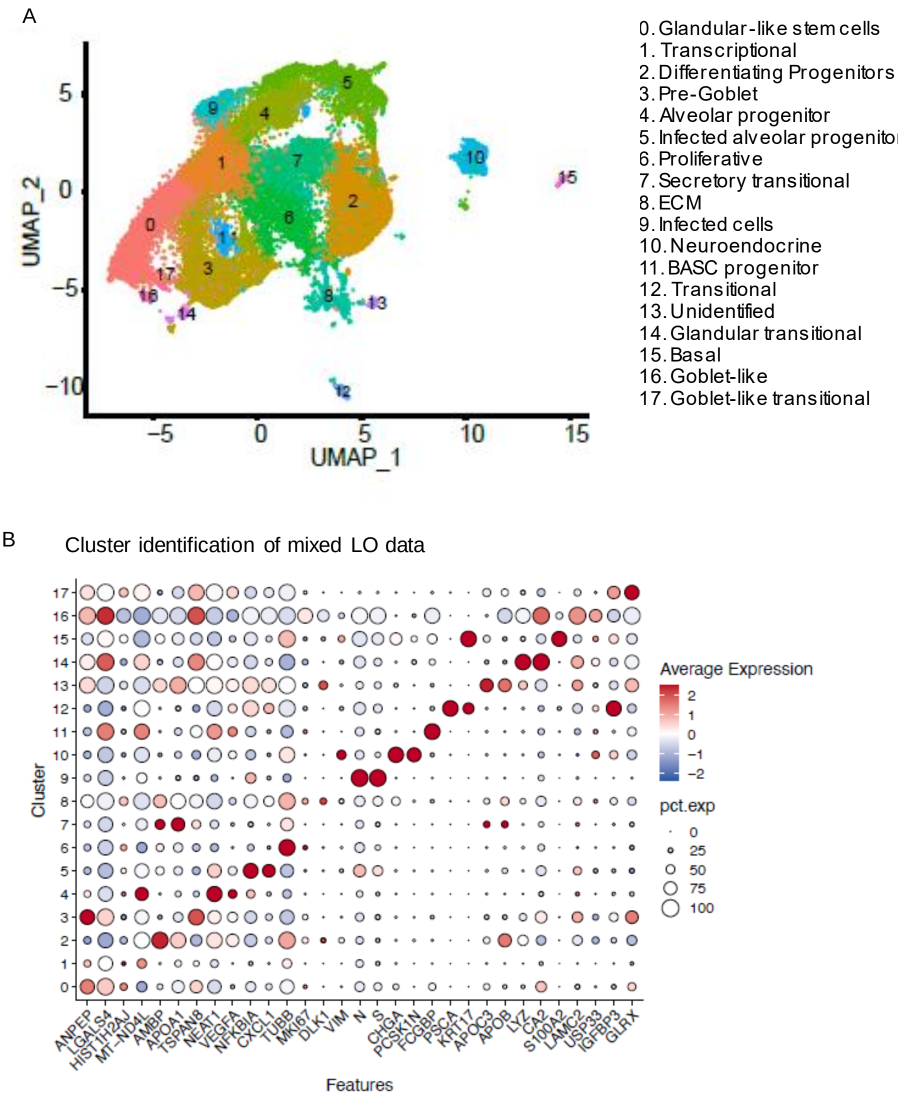

**FIGURE S6: Single cell transcriptional characteristics of the hPSC-LOs**. (A) Uniform manifold approximation and projection (UMAP) of hPSC-PLOs 16 hpi and (B) characteristic genes per cluster.

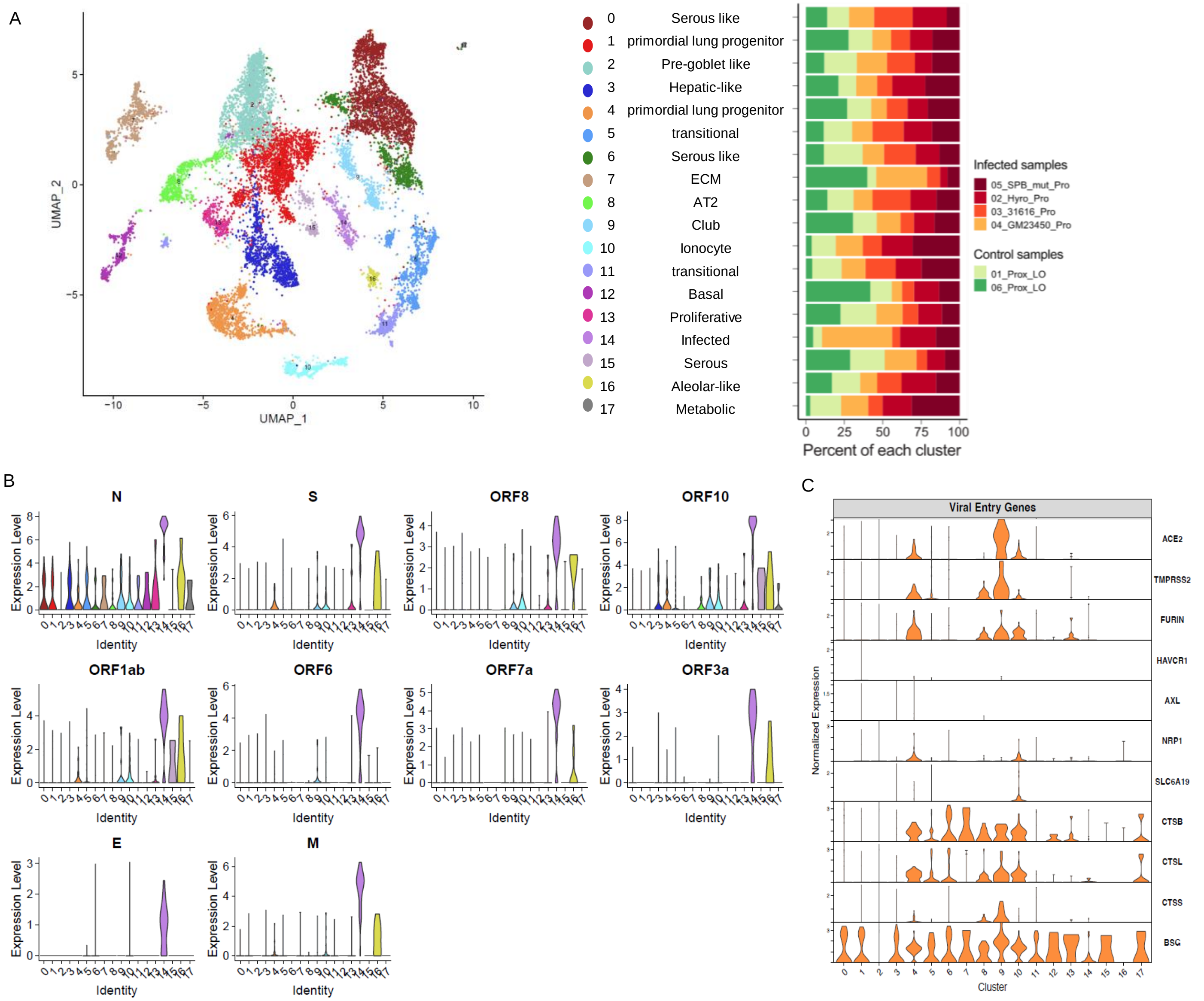

**FIGURE S7: Transcriptional changes and tropism in SP-B deficient hiPSC-LOs 16 hpi with SARS-CoV-2** (A) UMAP of 3D SP-B deficient PLOs, 24 hpi with the Alpha variant. Clusters were generated via Harmony Integration of 1 SP-B deficient PLO and 3 wild type PLO independent samples. ECM = extracellular matrix (B) SARS-CoV-2 viral genes in each cluster as violin plots. The vertical bars represent the expression level. (C) Violin plots of SARS-CoV-2 entry genes in each cluster in the PLOs. Most cells in the LOs get infected by SARS-CoV-2 with viral entry either via a canonical route (ACE2 and TMPRSS2) or a non-canonical route (e.g., endocytosis/macropinocytosis).

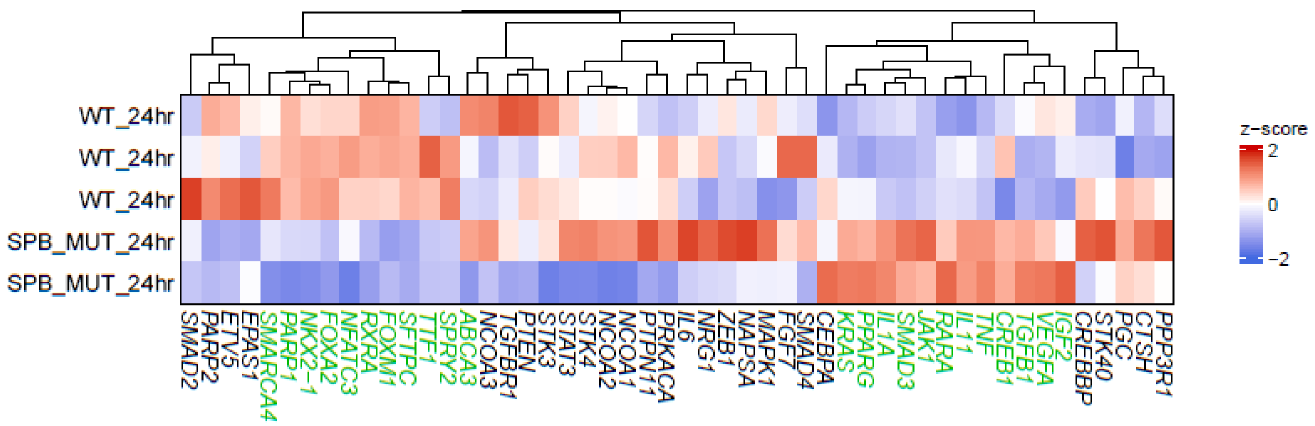

**FIGURE S8: Transcriptional changes in wild type (WT) and SP-B deficient PLOs 24 hpi.** Heat map showing average expression (represented as z-score) of key regulatory and transcriptional genes known to interact with surfactant protein B in WT and SP-B deficient 3D PLOs at 24 hpi with SARS-CoV-2. The genes differentially expressed between the infected wild type and SP-B deficient 3D PLOs are shown in green.

### Supplemental Information

### Supplementary Tables

**Supplementary Table 1:** Induced pluripotent stem cell lines

| Cell Name | Ethnicity | Sex | Age | Tissue | Produced Method | Type | Passage | Source |
|---|---|---|---|---|---|---|---|---|
| HYRO103 | Hispanic | Male | 31 | Hepatic fibroblast | Retroviral | iPSC | Not specified | ATCC |
| GM23450 | Black/African American | Female | 20 | Skin | Retroviral | iPSC | 29 | Coriell |
| GM23720 | White | Female | 22 | Peripheral vein | Episomal | iPSC | 44 | Coriell |
| Alda-31616 | White | Male | 25 | Lymphoblastoid Cell line | Episomal | iPSC | 10 | NIH |
| hiPro133 | White | Female | n/a | Skin | Retroviral | iPSC | 12 | CCRM |
| SP212-1-Cr3 Corr R11 | White | Female | n/a | Skin | Retroviral | iPSC | 20 | CReM |

iPSC= induced pluripotent stem cell

**Supplementary Table 2.** Profile of poractant alfa

| | **Portactant Alfa** |
|---|---|
| Source | Porcine |
| Phospholipid concentration (mg/mL) | 76 |
| Dipalmitoylphosphatidylcholine (mg/mL) | 30 |
| SP-B (mg/mL) | 0.45 |
| SP-C ($\mu$g/ protein/$\mu$mol $L^{-1}$ phospholipid) | 5.0-11.6 |
| Additives | No |
| Manufacturing | |
| Organic solvent extraction | Yes |
| Liquid-gel chromatography | Yes |

Adapted from https://curosurf.com/why-curosurf/surfactant-profiles/

**Supplementary Table 3:** Materials used in lung differentiations

| Name of Material/ Equipment | Company | Catalog Number |
|---|---|---|
| **Cell Culture** | | |
| 12 well plates | Corning | 3512 |
| 12-well inserts, 0.4um, translucent | VWR | 10769-208 |
| 2-mercaptoethanol | Sigma-Aldrich | M3148 |
| Accutase | Innovative Cell Tech | AT104 |
| ascorbic acid | Sigma | A4544 |
| B27 without retinoic acid | ThermoFisher | 12587010 |
| Bovine serum albumin (BSA) Fraction V, 7.5% solution | Gibco | 15260-037 |
| Dispase | StemCellTech | 7913 |
| DMEM/F12 | Gibco | 10565042 |
| FBS | Gibco | 10082139 |
| Glutamax | Life Technologies | 35050061 |
| Ham's F12 | Invitrogen | 11765-054 |
| HEPES | Gibco | 15630-080 |
| Iscove's Modified Dulbecco's Medium (IMDM) + Glutamax | Invitrogen | 31980030 |
| Knockout Serum Replacement (KSR) | Life Technologies | 10828028 |
| Matrigel | Corning | 354230 |
| Monothioglycerol | Sigma | M6145 |
| mTeSR plus Kit (10/case) | Stem Cell Tech | 5825 |
| N2 | ThermoFisher | 17502048 |
| NEAA | Life Technologies | 11140050 |
| Pen/strep | Lonza | 17-602F |
| ReleSR | Stem Cell Tech | 5872 |
| RPMI1640 + Glutamax | Life Technologies | 12633012 |
| TrypLE | Gibco | 12605-028 |
| Y-27632 (Rock Inhibitor) | R&D Systems | 1254/1 |
| **Growth Factors/Small Molecules** | | |
| Activin A | R&D Systems | 338-AC |
| All-trans retinoic acid (RA) | Sigma-Aldrich | R2625 |

| | | |
|---|---|---|
| BMP4 | R&D Systems | 314-BP/CF |
| Br-cAMP | Sigma-Aldrich | B5386 |
| CHIR99021 | Abcam | ab120890 |
| Dexamethasone | Sigma-Aldrich | D4902 |
| Dorsomorphin | R&D Systems | 3093 |
| EGF | R&D Systems | 236-EG |
| FGF10 | R&D Systems | 345-FG/CF |
| FGF7 | R&D Systems | 251-KG/CF |
| IBMX (3-Isobtyl-1-methylxanthine) | Sigma-Aldrich | I5879 |
| SB431542 | R&D Systems | 1614 |
| VEGF/PIGF | R&D Systems | 297-VP/CF |

**Supplementary Table 4:** Recipes of Medias used in lung differentiations

***3D organoid induction medium (day 17-22)***
Serum-free basal medium (see recipe) supplemented with:
FGF7 (10 ng/mL)
FGF10 (10 ng/mL)
CHIR99021 (3 µM)
EGF (10 ng/mL)

***3D organoid branching medium (day 23-28)***
*3D organoid induction medium* (see recipe) supplemented with:
All-trans retinoic acid (0.1 µM)
VEGF/PIGF (10 ng/mL)

***3D organoid maturation medium (day 29-34)***
3D organoid branching medium (see recipe) supplemented with:
Dexamethasone (50 nM)
Br-cAMP (100 µM)
IBMX (100 µM)

**3D Proximal lung organoid medium (day17-30)**
*Serum-free basal medium (SFBM)* supplemented with:
FGF10 (100ng/ml)
bFGF (250ng/ml)
Y27632 (10 µM)
Dexamethasone (50 nM)
Br-cAMP (100uM)
IBMX (100uM)

**3D Distal lung organoid medium (day17-30)**
*Distal- Serum-free basal medium (SFBM)* supplemented with:
CHIR99021 (3 µM)
SB431542 (10 µM)
FGF7 (10ng/ml)
RA (0.1uM)
Dexamethasone (50 nM)
Br-cAMP (100uM)
IBMX (100uM)

***AFE induction medium (day 4-6)***
Serum-free basal medium (see recipe) supplemented with:
SB431542 (10 µM)
Dorsomorphin (2 µM)

***DE induction medium (day 1-3)***
48.5 mL RPMI1640 + Glutamax
1 mL B27 without retinoic acid
500 µl HEPES (1%)
500 µl pen/strep
Human activin A (100 ng/mL)
CHIR99021 (5 µM) - only in the first 24 hours

***LPC induction medium (day 7-16)***
Serum-free basal medium (see recipe) supplemented with:
BMP4 (10 ng/mL)
All-trans retinoic acid (RA) (0.1 µM)
CHIR99021 (3 µM)

***Quenching medium 50ml***

49 mL DMEM/F12
1 mL FBS
500ul Pen/Strep

***Serum-free basal medium (SFBM) 500ml***
375 mL Iscove's Modified Dulbecco's Medium (IMDM) + Glutamax
125 mL Ham's F12
5 mL B27 without retinoic acid
2.5 mL N2
500 µl ascorbic acid, 50 mg/mL
19.5 µl monothioglycerol, 500 µg/mL
3.75 mL bovine serum albumin (BSA) Fraction V, 7.5% solution
5ml pen/strep

***Distal Lung Organoid Serum-free base medium (DLO-SFBM) 500ml***
475mL F12
16.65mL BSA
7.5mL HEPES
400µL $CaCl_2$
500µL ITS
5mL Pen/Strep
5ml B27 without retinoic acid

***Stem cell passaging medium (day 0) 50ml***
50 mL DMEM/F12 with Glutamax
13 mL Knockout serum replacement (KSR)
650 mL NEAA
130uL 2-mercaptoethanol
650uL pen/strep

**Supplementary Table 5.** SARS-CoV-2 strains

| Strain Name | Pango Lineage | Greek Name | Original sample sequencing links |
|---|---|---|---|
| USA/WA1-2020 | A | none "WA1" | GenBank: MN985325.1 |
| USA/NY-PV08410/2020 | B.1 | none "D614G" | GenBank: MT370900.1 and GISAID: EPI_ISL_421374 |
| hCoV-19/USA/CA_UCSD_5574/2020 | B.1.1.7 | Alpha | GISAID: EPI_ISL_751801 |
| hCoV-19/South Africa/KRISP-K005325/2020 | B.1.351 | Beta | GISAID: EPI_ISL_678615 with changes during passage at BEI listed in BEI datasheet https://www.beiresources.org/Catalog/animalviruses/NR-54009.aspx |
| hCoV-19/Japan/TY7-503/2021 | P.1 | Gamma | GISAID: EPI_ISL_792683 |
| hCoV-19/USA/PHC658/2021 | B.1.617.2 | Delta | none, changes from reference listed in BEI datasheet (https://www.beiresources.org/Catalog/animalviruses/NR-55611.aspx) and supplement to original publication (https://www.nejm.org/doi/suppl/10.1056/NEJMc2107799/suppl_file/nejmc2107799_appendix.pdf) |
| hCoV-19/USA/CA-SEARCH-59467/2021 | BA.1.20 | Omicron | GISAID: EPI_ISL_8186377 |

**Supplementary Table 6.** Antibodies

| Primary antibodies | Company | Catalogue # | Dilution rate |
|---|---|---|---|
| Nucleocapsid | GeneTex | GTX135357 | 1:1000 |
| Spike antibody | GeneTex | GTX632604 | 1:1000 |
| ACE2 | Cell signaling | 4355S | 1:500 |
| MUC5AC | Millipore | MAB2011 | 1:200 |
| SOX9 | R&D Systems | AF3075 | 1:200 |
| SCGB3A2 | abcam | ab181853 | 1:300 |
| HOPX | Santa Cruz Biotech | sc-398703 | 1:200 |
| proSPC | abcam | ab40871 | 1:250 |
| AcTub | | | 1:600 |
| SPC (mature) | LS Bio | LS-B9161 | 1:300 |
| SPB (mature) | 7 Hills | 48604 | 1: 500 (I) 1:500 (W)[a] |
| SPA | | | |
| SPD | | | |
| HTI-56 | Terrace Biotech | TB-29AHT1-56 | 1:150 |
| p63 | Boster | PB9152 | 1:250 |
| E-cadherin | | | |
| SMA | | | |

| Secondary Antibodies | Company | Catalogue # | Dilution rate |
|---|---|---|---|
| Donkey anti-mouse IgG (Alexa 488) | Abcam | Ab150105 | 1:500 |
| Donkey anti-mouse IgG (Alexa 555) | Abcam | Ab150106 | 1:500 |
| Donkey anti-mouse IgG (Alexa 647) | Abcam | Ab150107 | 1:500 |
| Donkey anti-rabbit IgG (Alexa 488) | Abcam | Ab150073 | 1:500 |
| Donkey anti-rabbit IgG (Alexa 555) | Abcam | Ab150074 | 1:500 |
| Donkey anti-rabbit IgG (Alexa 647) | Abcam | Ab150075 | 1:500 |
| Donkey anti-goat IgG (Alexa 488) | Abcam | Ab150129 | 1:500 |
| Donkey anti-goat IgG (Alexa 555) | Abcam | Ab150130 | 1:500 |
| Donkey anti-goat IgG (Alexa 647) | Abcam | Ab150131 | 1:500 |

### SUPPLEMENTARY VIDEOS

**Supplementary Video #1:** https://giphy.com/gifs/jTwljBoVJgmz0rFArn

**Supplementary Video #2:** https://giphy.com/gifs/eHEUP2X6fn4cPC5dp2

**Supplementary Video #3:** [https://giphy.com/gifs/ZBhrTM9f1j9FQqgXWq
